# Plasma titration provides a physical ruler for cross-platform proteomics

**DOI:** 10.64898/2026.08.09.740864

**Authors:** Yaqing Liu, Haiyan Wang, Yutong Zhang, Renquan Lu, Weihong Xu, Wanwan Hou, Yuyang Zhu, Xixi Li, Qingwang Chen, Yuechen Gao, Suhui Zhang, Qiaochu Chen, Zhiyin An, Jiaqi Yu, Wantao Ying, Xiaobo Yu, Jianhong Wu, Ziquan Fan, Xiao Yao, Zhihui Li, Hongmei Shan, Tiannan Guo, Zihong Ye, Xiaoping Yu, Gang Liu, Ying Yu, Zhiyu Peng, Chen Ding, Qiang Tian, Rui Zhang, Jinming Li, Xiang Fang, Fuchu He, Li Jin, Xingdong Chen, Leming Shi, Yuanting Zheng

## Abstract

Plasma proteomics is expanding across platforms and cohorts, and integrating these data for AI demands comparability at the protein level, not merely concordant associations^1,2^. Affinity and MS platforms use distinct probes (antibodies, aptamers, or peptides) and signal readouts, yielding contradictory cross-platform results^3,4^. Without a known quantitative truth, we cannot distinguish biology from measurement distortion, leaving no gold standard for integration. Here we introduce Plasmix—a plasma reference suite with predefined male:female ratios (M, 1:0; Y, 3:1; P, 1:1; X, 1:3; F, 0:1)—and show that preserving this quantitative titration gradient, not just technical repeatability, predicts cross-platform concordance and identifies protein measurements suitable for integration. Profiling Plasmix across five platforms (Olink, SomaScan, NULISA, AAgAtlas, and MS-DIA) and 12 protocols across 17 batches, we found discordance is dominated by signal generation, not sample identity, and platforms distort signals in a protein-specific manner. Crucially, proteins retaining the titration response showed stronger agreement in an independent cohort; anchoring to the Plasmix midpoint (P) via sample-to-reference ratios reduced distortions, extending harmonizable coverage by 10–20%. Plasmix thus provides a physical ruler to benchmark accuracy, identifying genuinely integrable measurements before pooling datasets or training AI models—a critical bottleneck for plasma proteomics.

## Introduction

Plasma proteomics is undergoing a rapid transformation. Population-scale affinity-based assays now measure thousands of circulating proteins in large cohorts, while recent mass-spectrometry (MS) workflows are extending both proteome depth and cohort scale^1,2,5–11^. These complementary advances are generating unprecedented datasets, yet they also present a growing challenge: how to integrate measurements generated by fundamentally different technologies. Cross-platform comparisons have revealed highly uneven agreement—protein-level correlations vary widely, and genetic, phenotypic, and disease associations can be shared, platform-specific, or even directionally discordant^3,4,12–18^. These inconsistencies undermine the confidence with which data from different measurement systems can be pooled, compared, or used as inputs for integrative analyses.

The implications of cross-platform discordance depend critically on the intended use of the data. For studies that analyse each platform separately, imperfect agreement may be tolerable so long as biological conclusions remain concordant^3,12,13,15,19^. Indeed, affinity–MS comparisons have shown that limited quantitative agreement between platforms among statistically significant proteins can coexist with concordant estimates of biological effects^4,20,21^. However, for direct data integration—pooling protein measurements across cohorts, comparing effect sizes between studies, transferring predictive models, or training AI algorithms—the field requires quantitative comparability at the protein expression level, not just agreement in downstream associations. Existing comparisons based on naturally varying cohorts can describe shared and platform-specific findings, but they lack a predefined, assay-independent expectation for the quantitative relationships among samples. They therefore cannot determine whether discordance reflects a simple change in scale or a fundamental distortion of relative sample differences, nor can they identify which measurements are genuinely integrable.

The sources of cross-platform discordance are multiple and deeply embedded in the measurement chain. Affinity assays and bottom-up MS derive signals from distinct molecular surrogates—antibodies, aptamers, peptides, or protein groups—that may represent different epitopes, modifications, or proteoforms of the same nominal protein target^4,13,16,20^. Background signal, detection limits, and normalization procedures further constrain effective measurement ranges and precision^17,22^. Sample matrix and pre-analytical handling can alter measured abundance in protein-and protocol-dependent ways^23–26^, while sample preparation workflows determine which proteins enter MS measurement, with depletion, enrichment, and neat-plasma approaches generating distinct proteome profiles^27–29^. In deep nanoparticle-based workflows, contamination from platelet, erythrocyte, and coagulation factors can inflate protein identifications and distort quantification^30^. Critically, these effects do not act uniformly across proteins; their net impact on any given measurement is difficult to predict without systematic benchmarking.

What the field has lacked is a physical benchmark—an experimental system with a known quantitative truth against which any measurement system can be assessed. Defined mixture samples have long been used to benchmark multi-laboratory quantitative performance across molecular profiling technologies^31–36^, but extending this design to intact plasma while preserving the complex matrix has remained a challenge. Such a system would enable prospective assessment of candidate platforms and protocols before large cohorts are profiled, and would provide an external criterion for evaluating whether data integration preserves the intended relationships among samples. It could also support common-reference anchoring, an emerging strategy for linking datasets across batches and platforms via the sample-to-reference ratio (SRR) method^37–39^. Recent recommendations for circulating blood proteomics have identified technology-agnostic reference samples as a means of linking diverse datasets^40^, but the benefits and limitations of this approach have not been systematically tested across plasma proteomic platforms, protocols, and study designs^41–44^.

Here we address this gap with Plasmix, a human plasma reference material suite comprising five samples with pre-defined male-to-female volumetric ratios (M, 1:0; Y, 3:1; P, 1:1; X, 1:3; F, 0:1). We profiled this titration series across five platforms—Olink, SomaScan, NULISA, AAgAtlas, and MS-DIA—encompassing 12 protocols and 17 analytical batches. The design allowed us to distinguish technical repeatability from accuracy (preservation of the known quantitative titration gradient), and to test whether measurements that retained this gradient showed greater concordance across batches, protocols, and platforms, as well as in an independent same-participant cohort. We examined how assay response, matrix context, and protein properties shaped quantitative distortion, and we evaluated reference-anchoring strategies under balanced and confounded study designs. Together, these analyses define the boundaries of cross-platform quantitative comparability, identify which protein measurements can be integrated with confidence, and establish a practical framework for benchmarking plasma proteomic data before they are pooled or used to train AI models. Plasmix thus provides a physical ruler for cross-platform plasma proteomics.

## Results

### Plasmix reveals fragmented coverage across the plasma proteome

Plasmix comprised five samples with pre-defined male-to-female volumetric ratios (M, 1:0; Y, 3:1; P, 1:1; X, 1:3; F, 0:1). NIST SRM 1950 (N) was co-measured as an independent plasma reference. The suite was profiled across five technology platforms and 12 protocols across 17 batches (**Fig. 1a**). Analytical features were harmonized to canonical protein identifiers while retaining distinct assay features targeting the same protein (**Supplementary Table 1**). This design enabled direct assessment of whether platform-specific measurements preserved predefined quantitative differences.

**Fig. 1.**
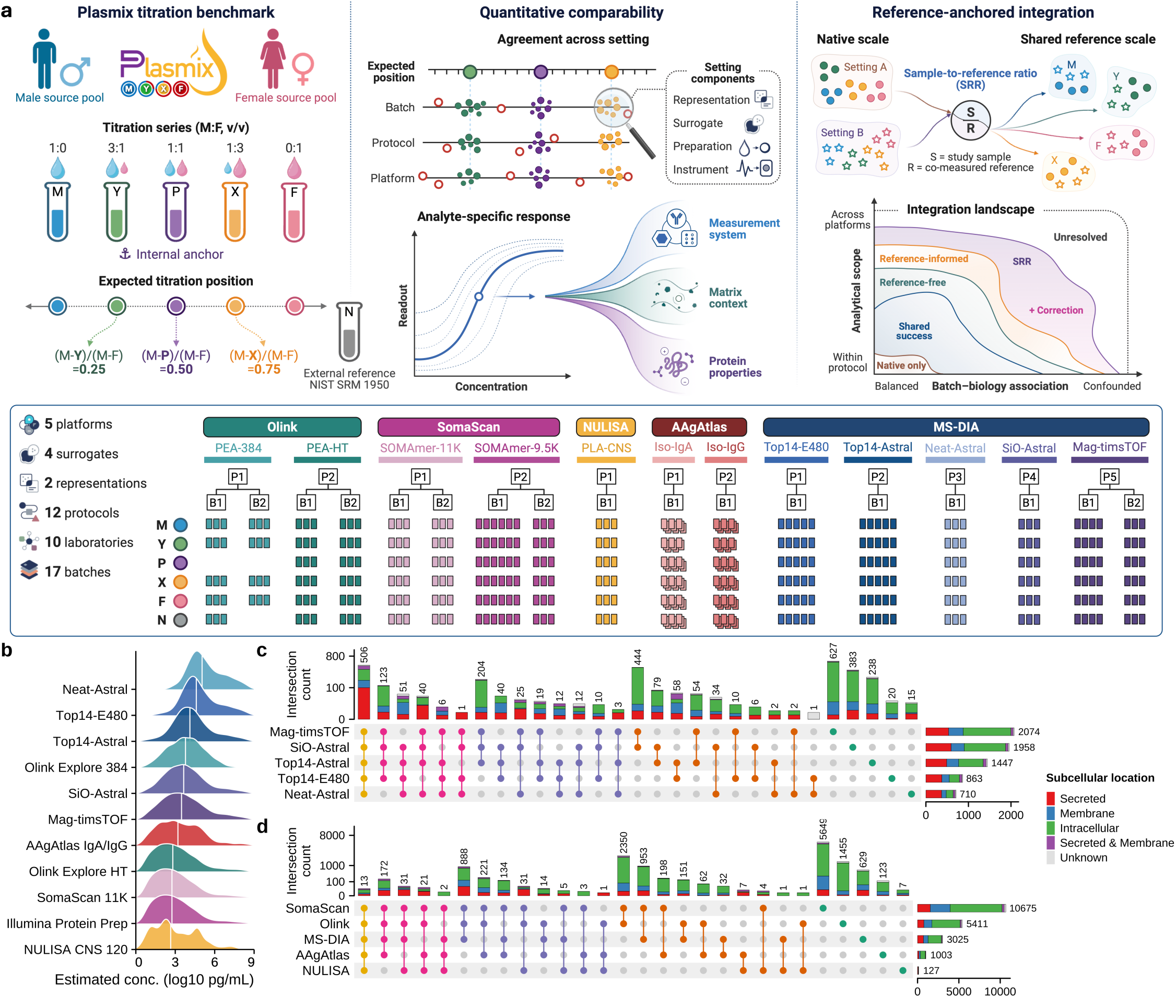
Plasmix study design and coverage of the plasma proteome. **a,** Schematic of the study design structured into five sequential phases to evaluate the boundaries and trade-offs of cross-platform integration. Plasmix reference materials, comprising sex-balanced, volumetrically titrated plasma pools (M, Y, P, X, F) alongside the NIST SRM 1950 (N) standard, establish biological and matrix ground truths. Titration benchmarks differentiate monotonic responses from distorted technical artifacts. Analyses then quantify integration bottlenecks driven by divergent representations, surrogates, and instruments. Next, the sample-to-reference ratio is evaluated for aligning batch effects. Finally, the framework delineates the analytical boundary where intrinsic physicochemical properties and kinetic limits constrain quantitative fidelity. The lower panel details the multi-center deployment across five platforms encompassing four molecular surrogates, two representation formats, and 12 distinct protocols spanning 17 batches. **b,** Ridge density plots illustrating the dynamic range of estimated protein concentrations across 11 specific analytical protocols. Baseline abundances (*log*_10_ pg/mL) are mapped from the Human Protein Atlas (HPA) reference database with proteins ordered by the median concentration stratum for each protocol. **c, d,** UpSet plots showing the intersection of detected proteomic features across five distinct DIA-MS pre-analytical workflows (**c**) and five major technology platforms including DIA-MS, SomaScan, Olink, NULISA, and AAgAtlas (**d**). Stacked bar charts within the UpSet plots indicate the corresponding sub-compartment distribution of the intersected features based on HPA subcellular localization annotations.

The five platforms and 12 protocols sampled remarkably distinct regions of the circulating proteome. Affinity-based assays extended broadly into low-abundance proteins, whereas neat-plasma DIA-MS was concentrated at higher abundance. Depletion and nanoparticle workflows shifted MS coverage towards lower abundance, yet their distributions remained distinct from those of the affinity platforms (**Fig. 1b** and **Supplementary Table 2**). Human Protein Atlas (HPA) blood-concentration annotations were available for 35.5–91.4% of protocol-level targets, and HPA-annotated proteins spanned most native measurement ranks within each protocol—indicating that concentration mapping was not confined to a narrow signal interval.

Despite the shared goal of quantifying plasma proteins, the platforms covered largely non-overlapping molecular territories (**Fig. 1c,d** and **Supplementary Table 3**). The five DIA-MS protocols shared only 506 proteins, whereas Mag-timsTOF alone measured 2,074 proteins, with 627 unique to that workflow. SomaScan provided the largest feature space (10,675 analysed features), of which 5,649 were not represented by any other platform. By contrast, only 888 features were shared among DIA-MS, SomaScan and Olink—the three high-throughput platforms most commonly used in population-scale studies.

This fragmentation extended to the biological nature of the detected proteins. SomaScan-exclusive targets and Mag-timsTOF-exclusive proteins were predominantly intracellular (59.4% and 68.0%, respectively), whereas shared features were enriched for secreted and membrane proteins. Thus, greater proteomic breadth came at the cost of fragmentation: each platform expanded coverage by accessing distinct molecular subsets, but this did not enlarge the shared quantitative measurement space required for cross-platform comparison and integration.

The Plasmix design (**Fig. 1a**) provided a physical bridge across these fragmented landscapes. By profiling the Plasmix alongside NIST SRM 1950 (N) across all platforms, we could now ask a question that platform coverage alone cannot answer: when platforms do measure the same protein, do their quantitative measurements agree—and if not, why?

### Repeatability does not guarantee quantitative accuracy

The titration benchmark immediately revealed a counter-intuitive principle: high technical repeatability does not ensure quantitative accuracy. We defined a protein measurement as preserving the expected response if it met two criteria across the M, Y, P, X, and F series: strict monotonic ordering (the intermediate samples fell in their correct sequence) and a titration response coefficient (TRC) deviating by less than 0.25 from the nominal positions (0.25, 0.50, and 0.75 for Y, P, and X, respectively). This TRC criterion—essentially a fidelity score—reports how closely each platform’s measurements track the known mixing ratios.

The ability to preserve the gradient depended primarily on the magnitude of the underlying biological difference (M/F). Measurements with strong M/F effects consistently passed the titration test, whereas weaker effects rarely did—regardless of platform or detection status (**Fig. 2a,b**). Yet even among the strongest effects, passage was not universal, indicating that a large signal does not automatically protect against distortion. Conversely, a minority of proteins with modest M/F differences still passed, suggesting that certain analytes are inherently more robust to platform-specific distortion.

**Fig. 2.**
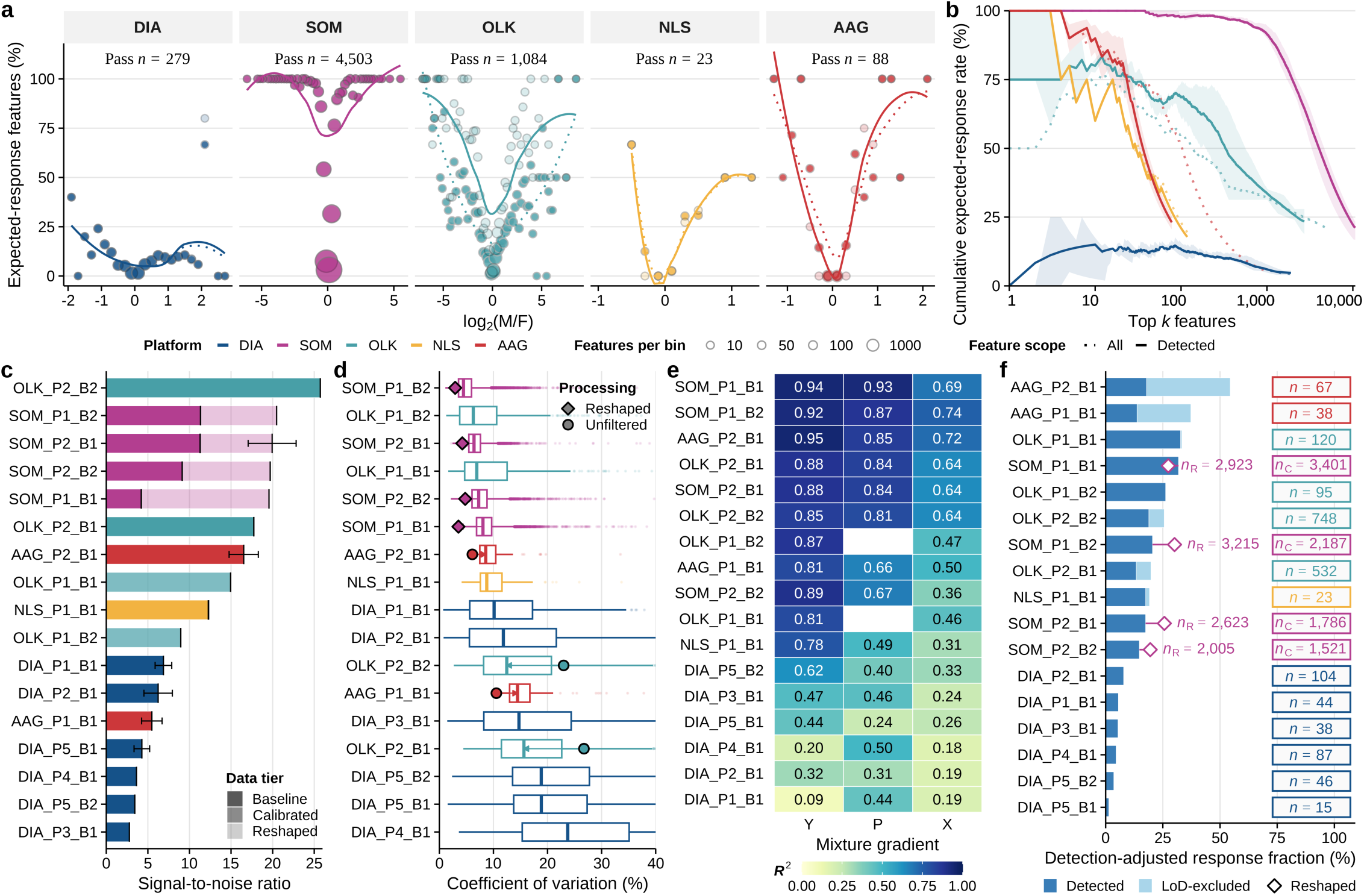
Benchmarking quantitative responses with the Plasmix titration series. **a,** Expected-response rates across the log₂(M/F) range. Feature–batch observations were binned at 0.2. Points show rates; size denotes observation count. Loess curves show all evaluable (dotted) and vendor-detected (solid) features. “Pass *n*” counts unique features passing in any primary-analysis batch, regardless of detection. **b,** Cumulative expected-response rates for features ranked within batches by descending absolute log₂(M/F). Lines show platform means for all evaluable (dotted) and vendor-detected features (solid); ribbons show their interquartile ranges across batches. **c,** PCA-based SNR for detected features across processing tiers. Bars show means across valid three-replicate combinations, error bars show s.d., and opacity denotes the data tier. **d,** Technical replicate CV distributions for vendor-detected features at the primary processing stage. Boxes show medians and interquartile ranges, with 1.5× interquartile-range whiskers. Symbols and arrows denote changes after detection filtering or SOMAmer-based reshaping. **e,** Gradient-fit *R*^2^ for Y, P and X mixtures among vendor-detected features at the primary processing stage. Cell values show fixed-prediction *R*^2^ by batch and mixture. **f,** Detection-adjusted expected-response fractions. Stacked bars distinguish vendor-detected from LoD-excluded expected-response features. Labels indicate the corresponding feature counts; open symbols show Reshaped SOMAmer-based values relative to Calibrated results. Expected response was defined as titration monotonicity and a mean absolute titration response coefficient (TRC) deviation of < 0.25 across at least two intermediate samples. Baseline, Calibrated and Reshaped denote basic correction, feature-wise inter-plate standardization and sample-specific normalization to an external reference, respectively.

Importantly, conventional quality metrics failed to predict this fidelity. Global sample separation (SNR) and feature-level repeatability (technical CV) captured different aspects of analytical performance, but neither reliably indicated whether a measurement preserved the known gradient (**Fig. 2c,d** and **Extended Data Fig. 1a–c**). A striking example: even among measurements within CV ≤10%, a substantial fraction (61.1–88.1%) of proteins failed the monotonicity criterion (**Extended Data Fig. 1c**). Gradient-fit *R*^2^ provided a complementary batch-level assessment of how closely each intermediate mixture followed its feature-specific expected response, revealing variation across platforms, batches and mixtures, with X generally showing lower fits than Y and P (**Fig. 2e** and **Supplementary Table 4**). In short, a measurement can be highly reproducible and still be quantitatively inaccurate.

The number of measurements meeting the TRC criterion varied widely across platforms, from 1.4% to 33.0% of vendor-detected features, with additional features falling below limit of detection contributing to the totals for Olink and AAgAtlas (**Fig. 2f** and **Supplementary Table 4**). Affinity-platform processing also altered the operationally detected feature space across stages: detection remained nearly complete in SomaScan microarray batches, increased markedly in sequencing-readout SOMAmer assays and decreased in Olink (**Extended Data Fig. 2a,b**). Beyond these changes in detection status, sample-specific normalization in SomaScan and IPP also affected quantitative-response preservation, shifting expected-response classifications in both directions and producing net gains in three batches and a net loss in one (**Extended Data Fig. 3a**). This reshaping also moved features across the statistical-significance threshold in both directions, showing that reshaping could not be assumed to improve quantitative fidelity uniformly (**Extended Data Fig. 3b,c**).

Together, these results establish that accuracy—preservation of a known quantitative relationship—is distinct from reproducibility and cannot be inferred from conventional quality metrics. The titration benchmark provides a direct, prospective way to identify which protein measurements can be trusted for cross-platform integration.

### Titration response predicts cross-platform reproducibility in human cohorts

If the titration response is a meaningful metric of measurement fidelity, then proteins that pass test should also show stronger agreement across platforms in real human populations. To test this, we first established an external benchmark of well-characterized sex-associated proteins. Using published data from UK Biobank, Iceland, Wellness, and BAMSE cohorts^3,45^, we classified proteins by the strength of evidence supporting a male–female difference. Tier 1 proteins—supported in at least three cohorts and by both Olink and SomaScan—showed a high level of agreement between the two independent study sources (*r* = 0.89; ρ = 0.92), whereas concordance declined progressively in lower evidence tiers (**Extended Data Fig. 4a–c** and **Supplementary Table 5**). These high-confidence proteins generally retained their direction in Plasmix, confirming the biological relevance of the M/F axis (**Extended Data Fig. 4d,e**).

Against this external evidence, the Plasmix gradient generated a substantially broader quantitative range than population cohorts. The central 95% width of the Plasmix Olink M/F distribution was 5.65 on the log₂ scale, compared with 0.20–0.76 across the four population cohorts (**Fig. 3a**). Among matched proteins, Tier 1 effects remained strongly correlated between UK Biobank and Plasmix (*r* = 0.88; **Extended Data Fig. 5a**). Even after matching the Plasmix donor numbers and mimicking sex-specific pooling, the UK Biobank M/F effect-size range remained substantially narrower (**Extended Data Fig. 5b**). Proteins showing little population-level sex difference but marked Plasmix shifts were enriched for intracellular protein complexes and nuclear components (**Extended Data Fig. 5c**), consistent with additional sample-context differences beyond population-averaged sex effects. The titration design thus acts as a stress test that amplifies biological contrasts to reveal platform-specific distortions that would otherwise remain invisible in population data.

**Fig. 3.**
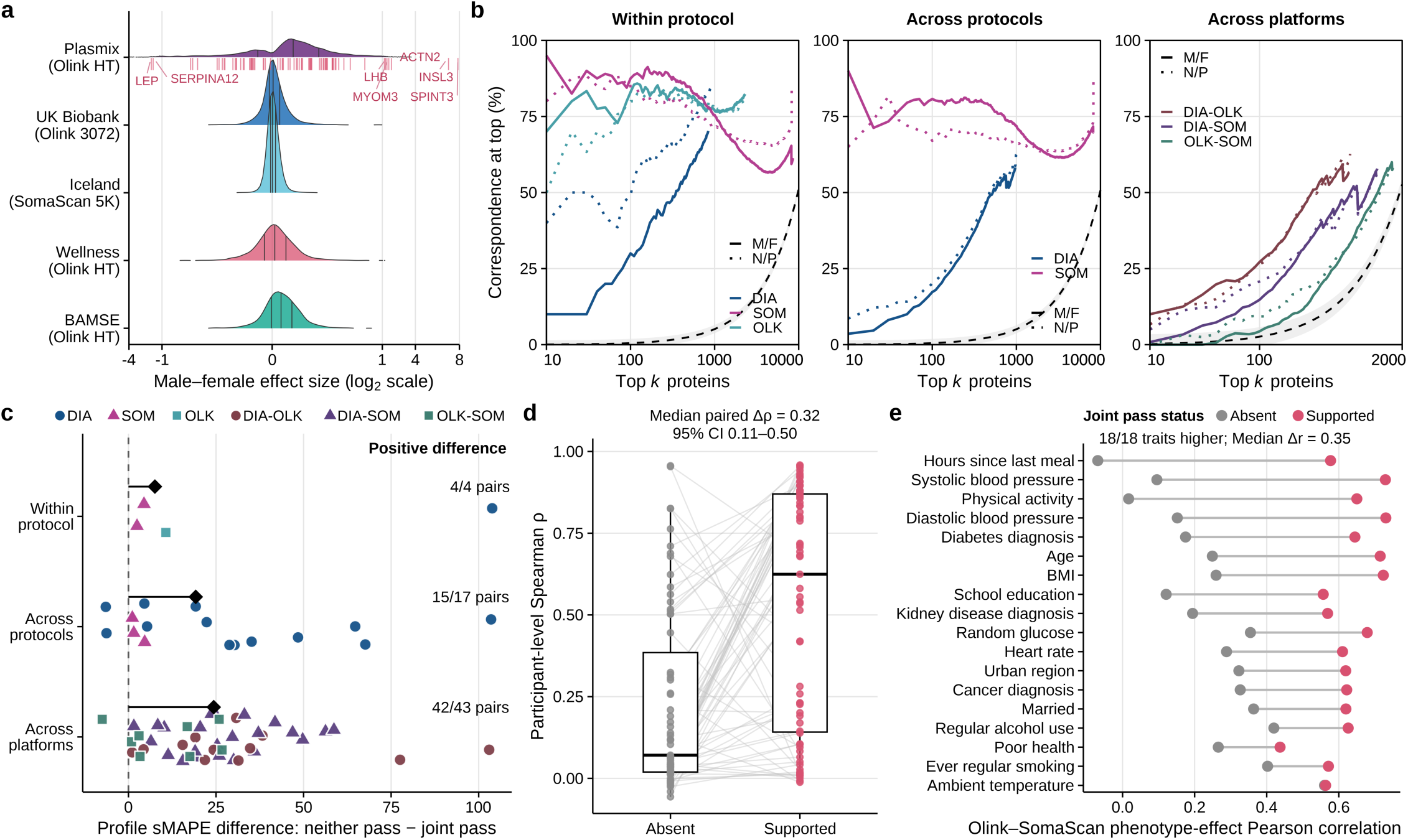
Quantitative concordance of protein measurements in relation to titration response. **a,** Distributions of male–female effect estimates in Plasmix and four population cohorts. Internal lines mark quartiles; pink ticks mark Tier 1 proteins. The x axis is compressed outside −1 to 1. **b,** Correspondence at the top (CAT) for proteins ranked by absolute M/F (solid) or N/P (dotted) effect across batch pairs within protocols, across protocols or across platforms. Colours denote platforms or platform pairs, and lines show means across batch pairs. Grey lines and ribbons indicate random-ranking expectations and 95% intervals. **c,** Batch-pair differences in median profile sMAPE (neither pass minus joint pass) after M/F-magnitude matching. Joint pass denotes expected-response passage in both batches and neither pass in neither; one-pass proteins were excluded. Profiles comprise M and shared intermediate samples relative to F. Positive values indicate lower joint-pass sMAPE. Points denote batch pairs, black diamonds denote medians and labels give positive/evaluated pair counts. **d,** Participant-level Olink–SomaScan Spearman correlations in China Kadoorie Biobank for Supported and M/F-magnitude-matched Absent proteins (*n* = 67 per group). Supported proteins had at least two jointly passing Plasmix Olink–SomaScan batch combinations; Absent proteins had none. Lines connect matched proteins; boxes show medians and interquartile ranges, with 1.5× interquartile-range whiskers. The annotation shows the median paired difference and 95% bootstrap confidence interval. **e,** Olink–SomaScan Pearson correlations of protein–phenotype effects across 18 non-sex traits in the same protein sets.

The broader effect range did not translate uniformly into transferable rankings. Although the N/P contrast showed greater quantitative separation and yielded more consensus differential signals than M/F, ranked-effect concordance remained protocol-and platform-dependent, and leading cross-platform overlaps sometimes approached random-ranking expectations (**Fig. 3b** and **Extended Data Fig. 6a–d**; **Supplementary Tables 6–8**). This prompted us to test whether preservation of the titration response could identify measurements with greater cross-setting concordance beyond differences in effect magnitude.

We next asked whether preserving the titration response predicted cross-platform agreement. For each batch pair, we classified proteins as jointly passing the titration test in both batches or neither passing. After matching the two groups for M/F effect magnitude, jointly passing proteins showed substantially lower cross-batch profile error (sMAPE) than neither-passing proteins in 61 of 64 evaluated batch pairs (**Fig. 3c**). The median advantage grew as analytical settings diverged: 7.6 percentage points between batches of the same protocol, 19.2 across protocols, and 24.3 across platforms. The titration criterion became increasingly informative as measurement conditions became more disparate.

This stratification generalized directly to independent population measurements. In the China Kadoorie Biobank^12^, 67 proteins with joint-pass support across Plasmix Olink–SomaScan batches had a median participant-level cross-platform correlation of 0.624. By contrast, effect-matched proteins without joint-pass support showed a median correlation of only 0.071—the median paired increase was 0.318 (95% bootstrap CI, 0.112–0.496; **Fig. 3d**). Moreover, joint-pass-supported proteins showed higher Olink–SomaScan correlations of protein–phenotype effect estimates for all 18 evaluated non-sex traits, with a median Pearson increase of 0.35 (**Fig. 3e**). These results demonstrate that the titration response, measured in a controlled laboratory benchmark, predicts whether a protein measurement will replicate across platforms in large-scale human studies—providing a prospective tool for prioritizing integrable proteins before cohorts are profiled.

### Assay response, matrix, and protein properties shape quantitative distortion

To examine why preservation of the same predefined titration gradient varied so much across analytical settings, we first characterized the empirical relationship between protein abundance and measured signal for each platform. These “response envelopes”—essentially, the transfer function between true and reported readout—differed markedly across platforms in signal span, shape, and background burden (**Fig. 4a,b** and **Supplementary Table 9**). DIA spanned approximately six log units of measured signal; Olink showed the steepest response slope (1.98); and SOMAmer-based assays had the greatest estimated background burden (0.34%). Within SOMAmer family, the microarray and sequencing readouts produced distinct envelopes. In DIA, magnetic-nanoparticle enrichment yielded a narrower apparent abundance span and 4.1–4.5-fold higher estimated background burden than silica-nanoparticle enrichment. Platforms thus transformed abundance into signal through fundamentally different response functions.

**Fig. 4.**
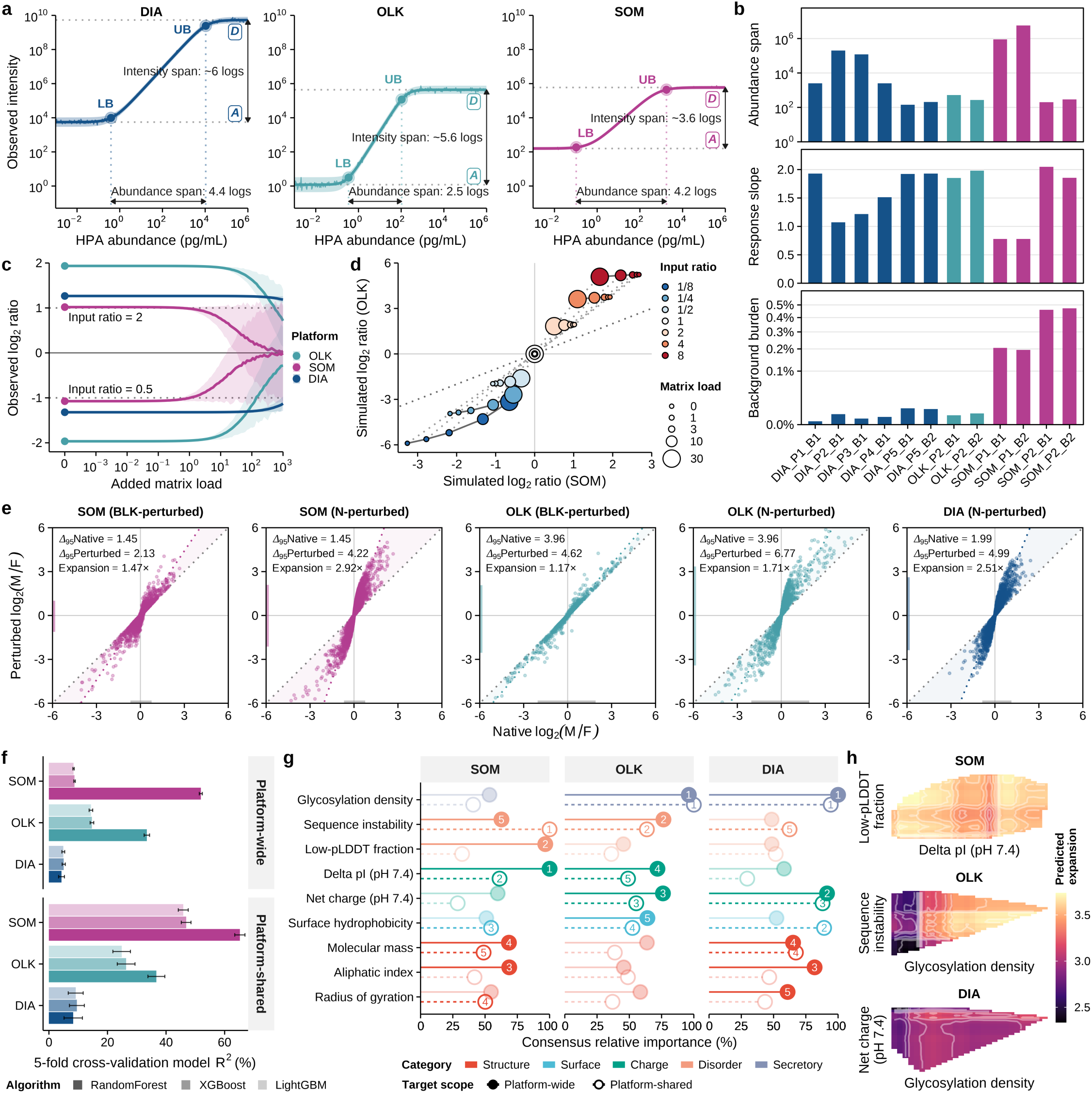
Assay response, matrix context and protein properties in quantitative distortion. **a**, Platform-level response envelopes relating HPA-derived blood abundance to measured intensity. Thick and thin lines show parameterized and stochastic trajectories; ribbons show variability. LB and UB mark response bounds; arrows indicate intensity and abundance spans. **b**, Batch-level abundance span, response slope and background burden. **c**, Simulated observed log₂ ratios for input ratios of 2 and 0.5 across matrix loads. Filled points mark zero load; lines and ribbons show means and 5th–95th percentiles. **d**, SomaScan–Olink projection of input ratios across matrix loads. Fill and size denote input ratio and matrix load; solid and dotted lines connect equal-ratio and equal-load conditions. **e**, Native and perturbed log₂(M/F) ratios after subtraction of NIST SRM 1950 (N) or buffer blank (BLK). Δ95 is the central 95% width, and expansion its perturbed-to-native ratio. **f**, Cross-validated R² for Random Forest, XGBoost and LightGBM models of log₂-transformed relative expansion, the N-subtracted log₂(M/F) ratio divided by its native value. Bars show mean ± s.d. across 50 repeated row-wise fivefold cross-validations of feature–batch observations. **g**, Mean normalized property importance across three algorithms for platform-wide and platform-shared targets. Displayed properties are the union of the platform-wide top five. Lines and symbols distinguish scopes; numbers mark top-five ranks within each scope. **h**, Two-dimensional partial-dependence surfaces from platform-shared Random Forest models. Axes show each platform’s two leading properties; fill denotes predicted relative expansion on the original scale.

These response differences were sufficient to generate systematic ratio distortion. When we projected fixed input abundance ratios (2 or 0.5) through the platform-specific envelopes, the observed log₂ ratios were already platform-dependent before added interference. Increasing matrix load—simulating background noise—progressively attenuated the ratios towards zero, with each platform following a distinct trajectory (**Fig. 4c**). Projecting the same input ratios through paired SomaScan and Olink envelopes produced curved deformation paths rather than a shared diagonal, meaning the same biological contrast would yield discordant quantitative ratios on different platforms (**Fig. 4d**).

Empirical matrix perturbations confirmed this prediction. Subtracting the NIST SRM 1950 reference plasma—a common “gold standard” material—from SOMAmer-based measurements expanded the M/F ratio distribution by 2.92-fold, compared with a 1.47-fold expansion from buffer-blank subtraction. NIST-plasma subtraction similarly expanded the ratios more than blank subtraction in Olink (1.71-fold versus 1.17-fold) and produced a 2.51-fold expansion in DIA (**Fig. 4e** and **Supplementary Table 9**). A single universal background correction factor cannot explain matrix-associated distortion; the effect is platform-and matrix-specific.

The susceptibility to perturbation also varied widely among proteins. Using a panel of 19 physicochemical properties (including charge, hydrophobicity, glycosylation, and structural disorder; **Supplementary Table 10**), we built machine-learning models to predict each protein’s distortion susceptibility. Across repeated cross-validation, Random Forest models explained 51.9% and 33.4% of the variance in SOMAmer-based and Olink analyses, respectively—rising to 65.2% and 36.7% for proteins measured across all platforms. By contrast, DIA models explained only 4.3%–8.3% of the variance (**Fig. 4f** and **Supplementary Table 11**). Thus, protein properties explain a substantial fraction of distortion in affinity-based assays, but very little in MS-based measurements.

The specific properties that mattered differed by platform. For SOMAmer-based assays, delta pI and low-pLDDT fraction (a measure of structural disorder) ranked highest. For Olink, glycosylation density and sequence instability were most important. For DIA, glycosylation density and net charge dominated (**Fig. 4g** and **Supplementary Table 11**). Two-dimensional partial-dependence surfaces revealed nonlinear, property-dependent response patterns, while substantial correlations among properties limited causal interpretation (**Fig. 4h**, **Extended Data Fig. 7a–d**, and **Supplementary Table 12**). Together, these analyses show that quantitative distortion emerges from the combined and interacting effects of assay response, matrix context, and protein properties—not any single dominant source. The mechanisms differ fundamentally between affinity and MS platforms, explaining why a one-size-fits-all correction is unlikely to succeed.

### A common reference anchor reduces platform-driven discordance

The mechanistic analyses revealed that distortion is platform-and protein-specific, suggesting that a uniform statistical correction is unlikely to succeed. But the quantitative landscape held a more striking revelation: the biological titration gradient was dwarfed by technical variation. Principal variance component analysis attributed 17.5% of total variation to measurement representation (signal intensity versus ratio), 12.8% to molecular surrogate (antibody, aptamer, or peptide), and 12.7% to their interaction. By contrast, the six plasma samples—the very biological differences we aimed to measure—accounted for only 0.02% of the total variance (**Fig. 5a** and **Supplementary Table 13**). The biological signal was embedded within vastly larger differences in how protein measurements were generated and reported.

**Fig. 5.**
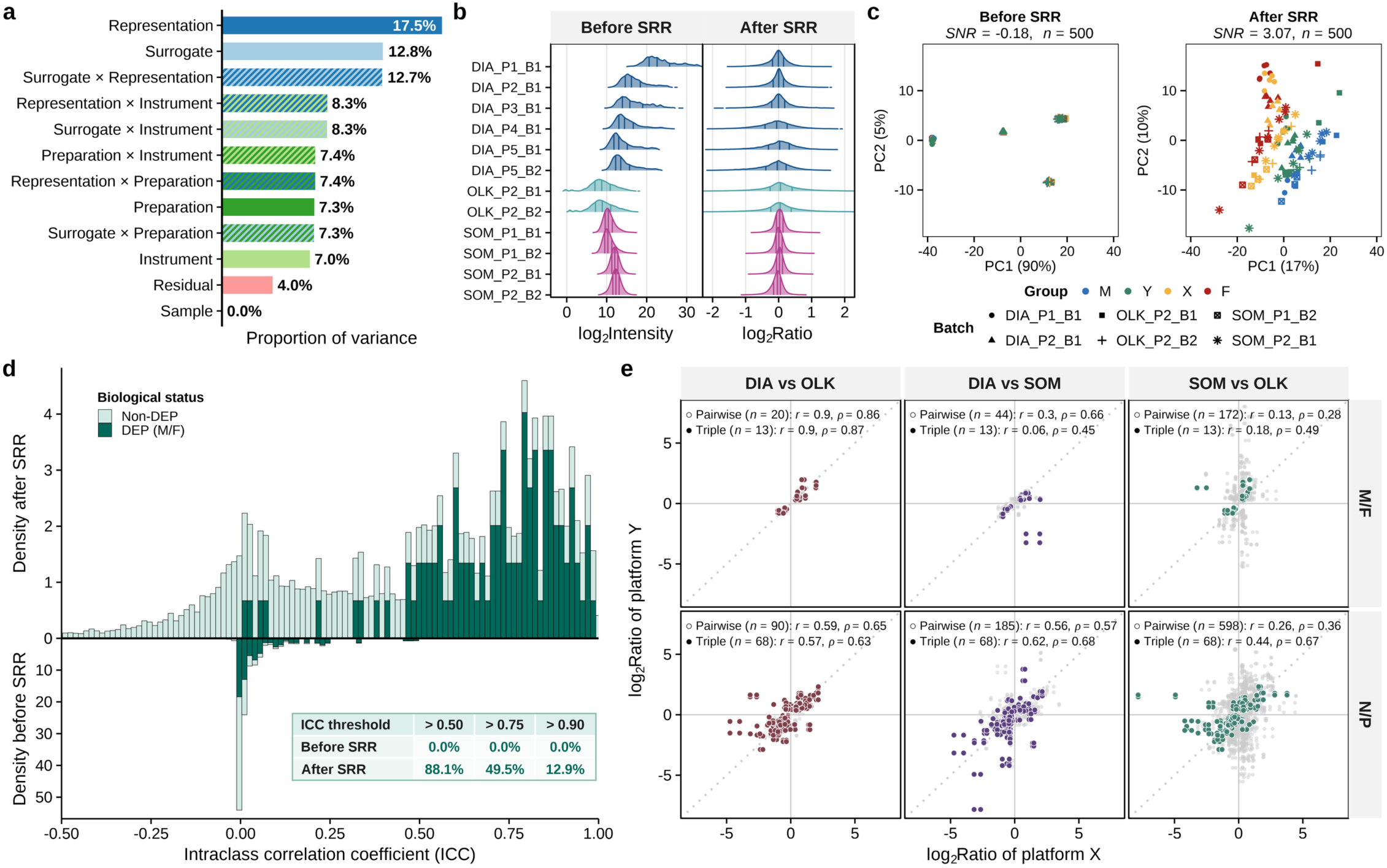
Effects of sample-to-reference ratios on quantitative discordance. **a,** Principal variance component analysis across the M, Y, P, X, F and N samples in 12 high-throughput batches. Bars show PCA-weighted variance proportions attributed to Sample, Residual and four technical factors: Surrogate (antibody, aptamer, or peptide), Instrument (scanner, sequencer or mass spectrometer), Representation (intensity or ratio) and Preparation (pre-instrument processing), together with pairwise interactions among the technical factors. **b,** Signal distributions by batch before and after sample-to-reference ratio (SRR) transformation. Ridge plots show log₂ intensities before SRR and log₂ sample-to-reference ratios after SRR; internal lines mark quartiles. **c,** PCA of six selected batches, comprising two each from DIA, Olink and SomaScan, before and after SRR transformation. Points are coloured by sample and shaped by batch; both panels use matched axis ranges. **d,** Protein-level intraclass correlation coefficients (ICCs) across the six selected batches before and after SRR transformation. ICCs were calculated across M, Y, X and F using a two-way agreement, single-measure model. Mirrored histograms are stratified by consensus M/F differentially expressed protein (DEP) status. The inset gives the percentages of consensus M/F DEPs with ICCs of at least 0.50, 0.75 and 0.90. **e,** Pairwise comparisons of log₂(M/F) and log₂(N/P) effects between platforms. Each point represents a feature in one cross-platform batch pairing. Grey points denote precision-verified DEPs shared by the corresponding platform pair; coloured points denote those shared across all three platforms. Annotations report the number of unique features, Pearson’s *r* and Spearman’s ρ.

This diagnosis pointed to a physical solution: rather than fitting a statistical model to the study samples alone, we could anchor every measurement to a concurrently measured common reference. We expressed each sample relative to the Plasmix midpoint P—a sample from the same source pools—by calculating sample-to-reference ratios (SRRs) within each batch and feature. This simple transformation required no model fitting and no assumptions about the distribution of the data.

The effect was dramatic. Native measurements occupied broad, platform-specific signal ranges that made cross-platform comparison nearly impossible. After SRR transformation, all measurements converged onto a common ratio scale centered at zero (**Fig. 5b**). At the sample level, platform and batch separation diminished, the M–Y–P–X–F ordering became clearer, and the signal-to-noise ratio increased from −0.18 to 3.07 (**Fig. 5c**). A shared denominator transformed a fragmented landscape into a coherent biological gradient.

SRR also improved protein-level reproducibility—especially for proteins carrying a genuine biological contrast. None of the 101 consensus M/F differentially expressed proteins reached an intraclass correlation coefficient (ICC) of 0.50 before SRR transformation. After transformation, 88.1% reached ICC ≥ 0.50, 49.5% reached ICC ≥ 0.75, and 12.9% reached ICC ≥ 0.90 (**Fig. 5d**). Among non-differential proteins, the proportion reaching ICC ≥ 0.50 increased more modestly (from 6.5% to 30.5%), indicating that SRR did not create uniform agreement by compressing all measurements; it selectively enhanced reproducible biological signals.

Despite this improvement, residual cross-platform differences remained. For M/F effects, Pearson correlations were 0.90 for DIA–Olink, 0.30 for DIA–SomaScan, and only 0.13 for SomaScan–Olink—with corresponding Spearman correlations of 0.86, 0.66, and 0.28 (**Fig. 5e**). Pearson correlations for N/P contrast were similarly modest (0.59, 0.56, and 0.26 for the same platform pairs). Restricting the analysis to proteins supported by all three platforms improved rank agreement in SomaScan-containing comparisons but did not eliminate off-diagonal measurements. Thus, SRR removed a major representation-dependent component of discordance, but protein-and platform-pair-specific differences persisted—raising the practical question of when and how reference anchoring can support integration in real-world study designs.

### Reference-anchoring defines where integration succeeds—and where it fails

The residual protein-and platform-pair-specific differences after SRR raised a practical question: can reference anchoring sustain integration when batch and biology were partially or fully confounded—the very scenario where statistical methods struggle most? To answer this, we designed a benchmark that paired 12 analytical batches into 66 batch pairs and evaluated each pair under three study designs: Balanced (both batches contain all samples), Partial (batches share only some samples), and Confounded (biological groups are separated across batches, with only the reference shared). Twenty integration pipelines were tested, including native data, five reference-free batch-effect correction algorithms, two reference-informed methods, SRR alone, and SRR followed by reference-free correction, producing 198 batch-pair–design tasks (**Fig. 6a**). Cross-batch identities of M, Y, X and F were hidden from all correction algorithms, mimicking real-world conditions where biological groups may be unevenly distributed. Harmonization was defined as joint achievement of cross-batch quantitative agreement and preservation of the expected response.

**Fig. 6.**
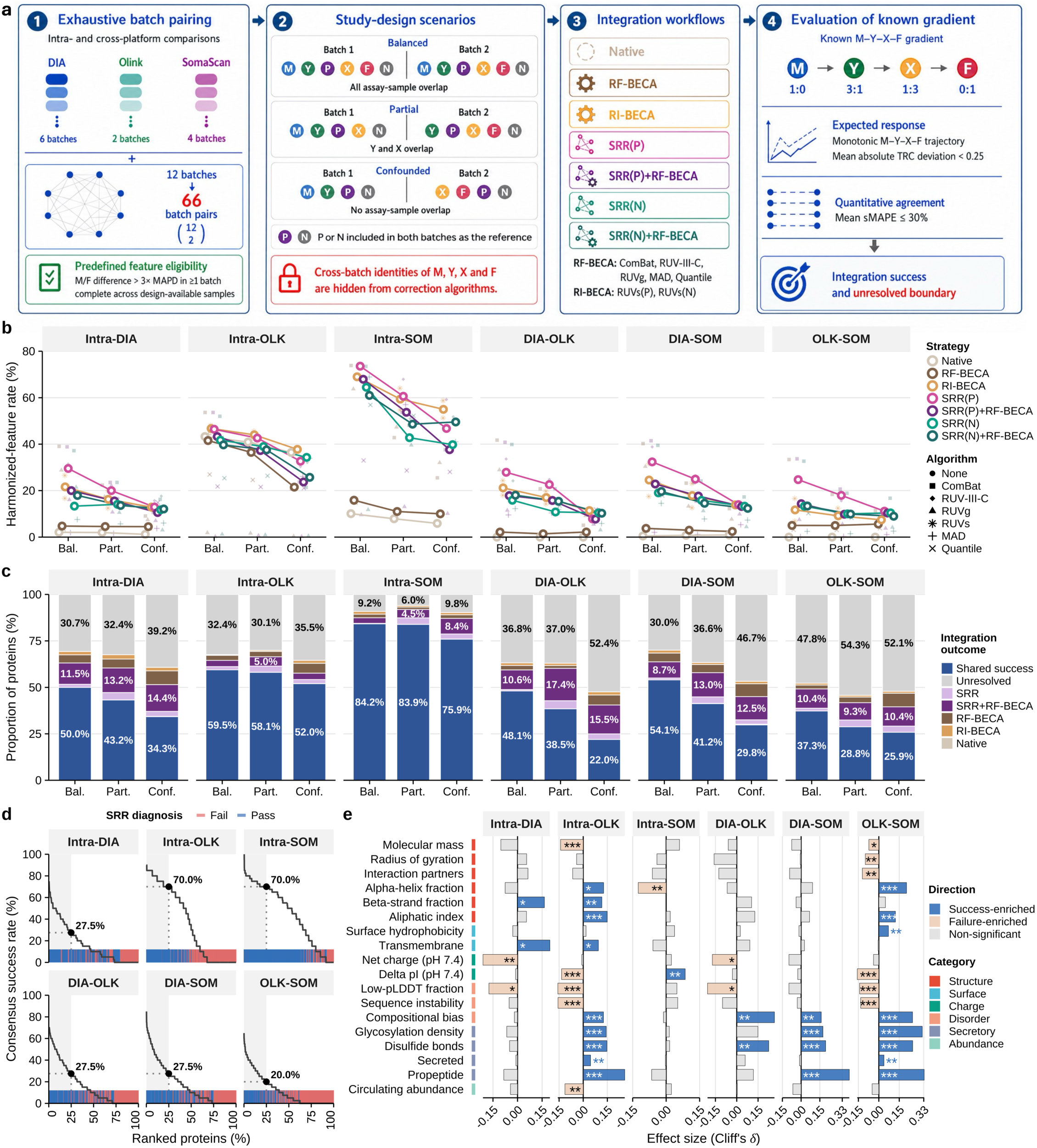
Integration performance across references, correction strategies and study designs. **a,** Schematic of the integration benchmark, including exhaustive pairing of 12 analytical batches into 66 batch pairs, Balanced, Partial and Confounded study-sample overlap designs, the 20 evaluated integration pipelines, and the criteria used to define harmonization. P or N was included in both batches as the selected reference, whereas cross-batch identities of M, Y, X and F were hidden from the correction algorithms. Quantitative agreement required mean sMAPE ≤ 30%; expected response required a monotonic M–Y–X–F trajectory and mean |TRC deviation| < 0.25; harmonized denotes joint passage of both criteria. **b,** Harmonized-feature rates across six within-and cross-platform scenarios and the three overlap designs. Small symbols denote individual algorithms after median summarization across batch pairs; coloured lines and open symbols denote medians for the corresponding strategy classes. **c,** Pair-averaged proportions of eligible proteins assigned to mutually exclusive integration outcomes under each overlap design: Shared success, Native, SRR, SRR plus reference-free batch-effect correction (SRR+RF-BECA), RF-BECA, reference-informed batch-effect correction (RI-BECA), or Unresolved. **d,** Consensus harmonization-success curves under the Balanced design. Proteins were ranked within each scenario by the median, across batch pairs, of the proportion of 20 pipelines satisfying the harmonization criterion. The rug indicates the corresponding SRR(P) pass or fail classification, and dotted guides mark the 25th ranked percentile. **e,** Cliff’s delta (δ) for physicochemical properties comparing upper-quartile consensus-success proteins with proteins showing 0% consensus success. Fill indicates enrichment direction; Two-sided Wilcoxon rank-sum test, \*\*\**P* < 0.001, \*\**P* < 0.01 and \**P* < 0.05.

Integration performance depended jointly on study design, platform pairing and correction strategy. At the strategy-class level, reference-enabled approaches—RI-BECA, SRR and SRR followed by RF-BECA—generally achieved higher median harmonized rates than Native or RF-BECA in Intra-SomaScan and cross-platform comparisons across the three study designs. Under Confounding, median rates were 37.6–55.0% for reference-enabled strategies in Intra-SomaScan and 7.4–14.0% across cross-platform settings, compared with 10.0% and 2.2–5.6%, respectively, for RF-BECA (**Fig. 6b** and **Supplementary Table 14**). This pattern was observed with both P-and N-based strategies, although their relative performance varied across settings. A co-measured reference therefore provided an explicit cross-batch coordinate when the study measurements alone could not distinguish batch effects from biology.

Reference anchoring also expanded the harmonizable protein set beyond what native measurements or unanchored correction alone could recover. Shared success—where both SRR-enabled and non-SRR methods succeeded—accounted for 84.2% of proteins in Balanced Intra-SomaScan comparisons and remained 75.9% under Confounding, though it was lower in cross-platform settings (**Fig. 6c** and **Supplementary Table 15**). Across the evaluated cross-platform settings, proteins recovered exclusively by SRR alone or by SRR followed by reference-free correction together accounted for an additional 9.7–21.8% of eligible proteins. Within this gain, SRR followed by reference-free correction contributed 3.0–17.4% of proteins not recovered by any other methods, whereas native-only success never exceeded 0.34%. These exclusive successes show that the reference supplied unique information for protein subsets that statistical correction alone could not recover.

Importantly, the benefit of reference anchoring was not confined to a reference constructed from the study sample pools, such as the Plasmix midpoint P. Under Confounding, P-and N-based strategies showed generally similar harmonized rates, with neither reference consistently superior across scenarios, indicating that an independently sourced plasma could also provide an effective cross-batch anchor for integration across studies (**Fig. 6b**). In Balanced comparisons with complete study-sample overlap, P yielded higher quantitative agreement in several settings, whereas expected-response retention was similar between P and N (**Extended Data Fig. 8a**). Even with P anchoring, INSL3, SPINT3 and LEP showed that reference effects remained protein-, platform-and batch-dependent (**Extended Data Fig. 8b**). Relative blank burden likewise varied across features and assay batches; subtracting it before SRR had little effect on expected-response retention but reduced quantitative agreement and harmonized rates across all three affinity-involving settings (**Extended Data Fig. 8c,d**).

Integration success varied continuously across proteins, ranging from passage across nearly all batch pairs and pipelines to zero success under the evaluated conditions, while SRR(P) successes were distributed throughout this continuum (**Fig. 6d**). The properties associated with success depended on the platform pair. DIA–Olink success favoured glycosylated and secreted proteins (Cliff’s δ = 0.27 and 0.23). DIA–SomaScan success was most strongly associated with higher circulating abundance (δ = 0.39). Olink–SomaScan success favoured higher abundance and disulfide-bonding proteins (δ = 0.33 and 0.32; **Fig. 6e** and **Supplementary Table 16**). Higher native measurement rank and HPA-estimated concentration also associated with success in several affinity-involving cross-platform settings, though these relationships were weak or reversed in some intra-platform analyses (**Extended Data Fig. 9a,b**). Reference anchoring expanded the integrable range, but platform pairing and protein context determined where that expansion was realized—and for a substantial fraction of proteins, integration remained unresolved.

## Discussion

We developed Plasmix—a plasma reference suite with a predefined titration gradient—to address a fundamental question in plasma proteomics: when are measurements from different platforms quantitatively comparable? By profiling this designed series across five platforms and twelve protocols, we showed that preserving a known quantitative gradient, rather than technical repeatability alone, predicts cross-platform concordance and identifies measurements suitable for integration. Common-reference anchoring via sample-to-reference ratios further reduced platform-driven discordance and extended harmonizable protein coverage. Yet integration remained protein-and platform-pair-specific, and a substantial fraction of proteins remained unresolved. Plasmix thus provides a physical ruler to benchmark accuracy, defining where plasma protein measurements are comparable, when reference anchoring extends integration, and which measurements remain beyond current capabilities.

The titration design adds a critical dimension that technical replicates and observational cohorts cannot supply: a known quantitative truth. Technical replicates establish precision; population cohorts reveal biological associations. Neither provides an independent expectation for how intermediate mixtures should behave. The Plasmix series therefore distinguishes technical repeatability from quantitative fidelity—and reveals that these two properties are surprisingly uncoupled. High precision did not guarantee gradient preservation, and platform-specific normalization that improved some measurements disrupted others. Accuracy cannot be inferred from reproducibility alone. This finding extends previous observations that participant-level measurements and downstream associations can exhibit different degrees of cross-platform concordance^3,12,15^, and it supports defined titrations as a prospective tool for prioritizing measurements likely to retain quantitative comparability before large cohorts are profiled.

The mechanistic analyses uncovered why fidelity varies across platforms and proteins. Quantitative distortion is not a single, removable batch offset; it reflects the combined effects of assay response, matrix context, and protein properties. Platforms transform abundance into signal through fundamentally different response functions, and these differences alone—before any added interference—can compress, expand, or reorder measurements of the same biological contrast^3,16,20,24,25^.Matrix perturbation further showed that interference is not a common additive floor. Protein properties explained a substantial component of distortion in affinity-based platforms but very little in MS-based measurements (**Fig. 4e–h** and **Extended Data Fig. 7**), indicating that the mechanisms of bias differ fundamentally between these technology families. Together with the dominant contributions of representation and molecular surrogate to total variance (**Fig. 5a**), these results define protein-and setting-specific limits to quantitative comparability and explain why a one-size-fits-all statistical correction cannot uniformly restore it.

Sample-to-reference ratios provided a physical solution to a physical problem. By expressing each measurement relative to a concurrently measured common reference, SRR reduced representation-dependent discordance without fitting a correction model to the study samples (**Fig. 5**). The transformation was simple, required no assumptions about data distribution, and selectively enhanced reproducible biological signals without compressing all measurements uniformly. A common denominator transformed a fragmented technical landscape into a coherent biological gradient. Under confounding between batch and biology—the scenario where statistical methods struggle most—SRR retained an explicit cross-batch coordinate that reference-free algorithms could not recover. Combining SRR with subsequent statistical correction further expanded the harmonizable protein set, but the gains depended on the algorithm and study design rather than accumulating uniformly (**Fig. 6c**).

The integration benchmark also revealed the boundaries of current capabilities. ComBat performed strongly in balanced designs, but its performance eroded under confounding. SRR excelled where study samples alone could not distinguish batch effects from biology, yet even optimal reference anchoring left a substantial fraction of proteins—approximately half of eligible targets in some cross-platform scenarios—unresolved. This is not a failure of the method; it is a fundamental characteristic of the measurements. Some proteins are inherently more robust to distortion, associated with higher circulating abundance, secretion, and specific physicochemical properties. Others are not. The field must therefore accept that quantitative comparability is not a binary property of a platform, but a continuous, protein-specific attribute that must be evaluated empirically for the intended platform pair, study design, and endpoint.

The implications for proteomic study design are clear. Shared identifiers define a common annotation space, but not a common quantitative space. Greater coverage does not automatically enlarge the protein space remaining comparable across settings (**Fig. 1c,d**). Defined mixtures can prospectively challenge candidate platforms and protocols, while references evaluated for the intended samples and procedures are particularly relevant when numerical pooling or transfer of quantitative estimates is planned. Comparability should be reported at the protein-measurement level for the specific platform pair and study design, rather than extrapolated to all shared targets.

The scope of these conclusions is defined by the material design and measurement systems examined. Plasmix specifies mixture ratios and predefined intermediate positions rather than traceable protein-specific concentrations, and stringent titration assessment is restricted to proteins with sufficient endpoint response. The material represents a single biological axis in pooled plasma from healthy Chinese donors, and the evaluated protein space reflects both the material and preparation workflows tested rather than the complete plasma proteome^46,47^. The conclusions are also conditional on the platforms, protocols, and processing generations examined. Extending this strategy with orthogonal biological contrasts, additional plasma backgrounds, pre-analytical conditions, and independently manufactured references could progressively enlarge the protein space evaluated under predefined quantitative constraints.

Despite these limitations, Plasmix demonstrates a principle that extends beyond this specific material: how experimentally defined plasma gradients can test comparability and reference performance before large plasma-proteomic datasets are pooled or quantitative results are transferred across measurement systems. As plasma proteomics moves towards clinical and regulatory applications, the need for such physical benchmarks will only grow. Plasmix provides a starting point—a physical ruler for cross-platform plasma proteomics—and a framework for building the broader coordinate system that the field will need to realize the promise of integrative, AI-driven discovery.

## Methods

### Reference materials

The study protocol was approved by the Ethics Committee of the School of Life Sciences at Fudan University (Approval No. FE253741). We recruited 110 healthy Chinese volunteers (55 men and 55 women; 20–60 years) without known disease or medication use during the preceding week; women were not menstruating at collection. After an overnight fast, blood was collected into EDTA-K2 tubes and plasma was separated within 4 h by centrifugation at 3,000 × g for 15 min followed by 16,000 × g for 10 min. Plasma was obtained from 108 participants. Donations were screened for HBsAg, HCV, Treponema pallidum and HIV, and biological sex was verified by SRY qPCR of cell-free DNA. One donation failed pathogen screening and two failed SRY verification, leaving 105 qualified donors (54 men and 51 women) and 1,790 ml and 1,690 ml of male and female plasma, respectively. Qualified donations were pooled by sex to generate male (M) and female (F) pools. Intermediate materials Y, P and X were prepared at M:F volumetric ratios of 3:1, 1:1 and 1:3, respectively. All five Plasmix materials were aliquoted, stored at −80 °C and verified by SRY qPCR.

NIST Standard Reference Material® 1950 (Metabolites in Frozen Human Plasma) was included as a heterologous plasma reference^48^. It comprises lithium-heparin plasma pooled from 100 fasted healthy US donors (50 men and 50 women; 40–50 years), processed within 60 min, centrifuged at 8,000 × g for 25 min, aliquoted and stored at −80 °C.

### Proteomic data generation and harmonization

Protein profiles were generated using data-independent-acquisition mass spectrometry (DIA-MS), Olink proximity extension assays, SOMAmer-based SomaScan and Illumina Protein Prep (IPP) assays, NULISAseq and the AAgAtlas autoantibody microarray. Olink, NULISA and SOMAmer-based samples were randomized after fixed calibrator, quality-control and blank positions were assigned. DIA-MS samples were analysed in repeated M–Y–P–X–F–N loops with non-consecutive technical replicates. AAgAtlas used three independent tubes per sample, each analysed in consecutive technical triplicates; one Y replicate that failed quality control was excluded. Detailed wet-laboratory procedures, run orders and batch-specific settings are provided in the Supplementary Methods and Supplementary Table 17.

DIA-MS comprised high-abundance-protein depletion, neat-plasma digestion, nanoparticle enrichment^49^ and magnetic-bead enrichment^50^ on Orbitrap Exploris 480, Orbitrap Astral and timsTOF Pro 2 instruments. Participating laboratories used locally optimized library-based or direct-DIA workflows and returned processed protein-expression matrices. Olink Explore 384 Cardiometabolic and Explore HT measurements were normalized to block-specific extension controls and then feature-wise to plate controls, producing ExtNPX and NPX, respectively^51^. SomaScan 11K Assay v5.0 measurements underwent hybridization normalization, calibrator median normalization, plate scaling, feature-specific calibration and Adaptive Normalization by Maximum Likelihood (ANML). IPP 9.5K used the same SOMAmer chemistry with a sequencing readout and produced ReadoutNorm, PlateNorm and SampleNorm outputs. NULISA CNS Disease Panel 120 counts were normalized sequentially to internal and inter-plate controls and converted to NPQ^52^. AAgAtlas IgG and IgA signals were the median fluorescence intensities at 532 nm and 635 nm, respectively; signal-to-noise ratio (SNR) was the median spot intensity divided by the 25th percentile of negative-control intensities^53^. This spot-to-background AAgAtlas SNR was distinct from the PCA-based sample-separation SNR used in the titration benchmark. Manufacturer quality-control criteria were applied to affinity-platform outputs.

Platform-reported assay identifiers were treated as analytical features, including SOMAmer probes, antibody pairs, array antigens and MS protein groups. Features were mapped to canonical UniProt accessions and classified as one feature to one protein, multiple features independently targeting one protein or one feature mapping to multiple proteins. Multiple features targeting the same protein were retained separately because they may represent distinct epitopes, modifications or peptides. Unmapped and one-to-many features were excluded, yielding 13,159 harmonized features. Platform outputs were organized into four tiers: Readout (raw counts or fluorescence); Baseline (detector-level output after basic internal-control or hybridization correction); Calibrated (feature-wise inter-plate standardized output); and Reshaped (SomaScan ANML or IPP SampleNorm, which applies sample-specific normalization against an external population reference). Canonical UniProt accessions were also used to map features to Human Protein Atlas blood concentrations and protein-class annotations^45^. When multiple concentration records were available, the median was used, prioritizing MS over immunoassay estimates; values were converted to pg ml⁻¹ and log₁₀-transformed.

Detection was evaluated at the Baseline tier for DIA-MS and AAgAtlas and at the Calibrated tier for Olink, NULISA and SOMAmer-based assays. Detection required AAgAtlas SNR ≥3, a non-zero finite DIA-MS intensity, or a platform-specific threshold derived from buffer blanks, negative controls or fixed background counts. Within a batch, a feature was detected in a sample group when more than half of the structurally available technical replicates passed and was detected overall when this occurred in at least one study or reference group. For titration-response, PCA-based SNR and technical-CV analyses, detection was defined across M, Y, P, X and F only. For differential analyses, missing values were imputed within batch, tier and processing stage from a Gaussian distribution centred on the 0.1th percentile of finite intensities with s.d. equal to 5% of its absolute value and scale-appropriate lower bounds. Detection and eligibility were determined before imputation. Global analyses retained features detected in >50% of batches and used K-nearest-neighbour imputation (k=10); PCA required detection in >80% of batches and stable detection in P^54^. PVCA used limit-of-detection imputation for complete batch absence followed by K-nearest-neighbour imputation for scattered missing values.

### Titration benchmarking

Primary titration analyses used Baseline-tier DIA-MS and AAgAtlas measurements and Calibrated-tier Olink, SomaScan and NULISA measurements; SomaScan Reshaped outputs were retained for processing-stage comparisons. To standardize batches with different replicate depths while preserving acquisition-round structure, all distinct combinations of three acquisition rounds shared across eligible sample groups were enumerated. Within each combination, the mean finite log₂ measurement defined a group centre when at least two selected measurements were available; no missing values were imputed. The linear-scale centre and the titration response coefficient for intermediate sample S were defined as shown below. This fixed M and F at 0 and 1 and assigned nominal positions of 0.25, 0.50 and 0.75 to Y, P and X. At least two finite intermediate TRCs were required. The mean absolute TRC deviation was the equally weighted mean of available intermediate-specific deviations and passed at <0.25. Adjacent relations among evaluable samples were supported when the earlier TRC was strictly smaller than the later TRC; ties failed. A feature passed monotonicity when at least two adjacent relations were defined, every relation was evaluable in at least one replicate combination and every relation was supported in >50% of evaluable replicate combinations. Expected response required both monotonicity and the TRC-deviation criterion; detection was reported separately.

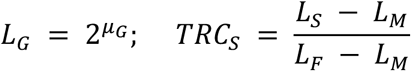

Batch-level gradient fit compared each observed Y, P or X response relative to F with the response expected from linear mixing on the abundance scale, without fitting an intercept or slope. PCA-based SNR quantified separation among available Plasmix groups relative to technical-replicate dispersion using variance-weighted distances in the first two principal components, following the ratio-based benchmarking framework used for Quartet materials^38^. Technical CV was calculated on the linear scale for complete triplicate groups and averaged across eligible groups and replicate combinations. Cross-batch quantitative agreement was calculated from the two batches’ arithmetic mean values for each evaluation sample and then averaged across the design-specific shared or validation samples.

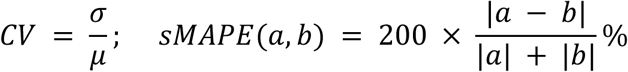

### Cross-setting validation

Correspondence-at-the-top analysis compared proteins ranked by absolute M/F or N/P effect across 12 high-throughput batches^55^. At each top-ranked fraction, correspondence was the proportion simultaneously present in both lists with concordant direction; random expectations used observed direction frequencies in the common-protein universe. For profile comparisons, proteins were classified as joint pass, one pass or neither pass for each batch pair. Joint-pass and neither-pass proteins were matched without replacement by Hungarian optimization of log₁p-transformed mean M/F effect magnitude. Profile agreement was calculated as sMAPE across M and at least two shared intermediate samples after expression relative to F.

Independent validation used published China Kadoorie Biobank Olink–SomaScan data^12^. Strict one-to-one assay mappings were retained. Plasmix support was defined from two Olink Explore HT and two non-ANML SomaScan batches: Supported proteins had at least two jointly passing Olink–SomaScan batch combinations, whereas Absent proteins had none. Groups were matched 1:1 by Plasmix M/F effect magnitude. Participant-level correlations were compared within pairs, with percentile 95% confidence intervals from 1,000 bootstrap resamples; protein–phenotype effect correlations were evaluated across 18 non-sex traits. Sex-associated proteins from UK Biobank, Iceland, Wellness and BAMSE were harmonized to the M/F direction and assigned evidence tiers according to cohort and platform support^3,45^. Plasmix Olink M/F effects were additionally compared with individual-level UK Biobank data. In 5,000 iterations, 54 men and 51 women were sampled without replacement, and direct and pseudo-pooled effects were summarized by their central 95% widths.

### Differential expression and consensus analyses

Differential expression was analysed with limma separately within each batch, tier and processing stage^56^. Exactly three technical replicates were selected once for each eligible sample group using a fixed random seed and retained across stages and contrasts. Features detected in at least one contrasted group were retained, missing values were imputed as above, moderated statistics were obtained by empirical Bayes and P values were adjusted by the Benjamini–Hochberg procedure^57^. Feature-specific technical variation was summarized on the log₂ scale as the median absolute pairwise difference (MAPD) among technical replicates. Verified differentially expressed proteins met all three criteria shown below; the remaining results were classified as non-significant, precision-rejected or small-magnitude as defined in the Supplementary Methods.

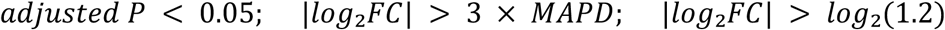

Consensus differential proteins were identified from Baseline-tier DIA-MS and Calibrated-tier Olink, SomaScan and NULISA data; AAgAtlas was excluded because it measured autoantibody binding rather than circulating protein abundance. A candidate direction required support from at least two platforms and four analytical observations. Effects were summarized by the median log₂ fold change, P values were combined by Fisher’s method and adjusted by Benjamini–Hochberg. Consensus proteins required a Benjamini–Hochberg-adjusted P value <0.05 and |median log₂ fold change| >log₂(1.2). Gene Ontology over-representation analyses used clusterProfiler and org.Hs.eg.db with all harmonized measured proteins having valid mappings as the shared background; terms were simplified at a semantic-similarity cutoff of 0.65^58^.

### Mechanisms of quantitative distortion

Principal variance component analysis (PVCA) was performed on 12 high-throughput DIA-MS, Olink Explore HT and SomaScan batches using Baseline-tier DIA-MS and Calibrated-tier affinity measurements. Unit-variance-scaled principal components explaining 80% of cumulative variance were entered into Bayesian mixed-effects models implemented in brms^59^. Models included five main effects—Surrogate, Representation, Preparation, Instrument and Sample—and pairwise interactions among the four technical factors. Weakly informative Student-t priors were used, and variance components were eigenvalue-weighted and aggregated across components.

Batch-and platform-level empirical response envelopes were constructed from Baseline-tier measurements converted to the linear scale. Feature–sample observations from M, Y, P, X, F and N were divided into 102 consecutive native-intensity intervals and summarized by mean signal, mean Human Protein Atlas blood abundance, median replicate CV and abundance-annotation coverage. A four-parameter logistic envelope was parameterized from empirical lower and upper signal anchors, an abundance midpoint and the abundance interval spanning 1–99% of modeled signal amplitude. Background burden was estimated from inner-tail intervals, and lower-, middle-and upper-range CVs generated variability envelopes. Within-platform simulations projected abundance ratios of 2 and 0.5 through each envelope over increasing dimensionless matrix load, using 1,000 simulations per platform, ratio and load. Cross-platform simulations projected input log₂ abundance differences from −3 to 3 through paired SomaScan and Olink envelopes at matrix loads of 0, 1, 3, 10 and 30. Full equations are provided in the Supplementary Methods.

Matrix perturbation was evaluated from detected Baseline-tier measurements on the linear scale. Technical replicates were summarized by their median intensities. Native M/F ratios were compared with ratios after subtraction of the feature-and batch-specific buffer-blank signal for affinity assays and after subtraction of NIST SRM 1950 plasma. DIA-MS was not evaluated under blank subtraction because corresponding blanks were unavailable. Distributional distortion was summarized by the central 95% width and its expansion relative to the native distribution. Feature-level susceptibility to NIST-plasma perturbation was the log₂-transformed relative expansion of the perturbed ratio versus the native ratio; near-zero native ratios and non-finite or non-positive relative expansions were excluded.

Thirty candidate structural and physicochemical properties were compiled from sequence, structure, annotation and reference-abundance resources. Correlation-and interpretation-guided filtering retained 22 non-redundant properties; circulating abundance, α-helix fraction and β-sheet fraction were excluded from prediction because their missingness was associated with expansion, leaving 19 predictors. Random Forest, XGBoost and LightGBM regressors were trained separately for DIA, Olink and SOMAmer-based assays in platform-wide target sets and in platform-shared target sets restricted to canonical proteins represented in all three modelled platform families^60–62^. Random Forest used 500 trees and terminal-node size 5; XGBoost and LightGBM used 100 boosting rounds, learning rate 0.05 and shallow trees. Performance was assessed using 50 repeats of row-wise fivefold cross-validation with identical folds across algorithms. Final models fitted to all eligible observations were used for normalized importance, partial-dependence and accumulated-local-effect profiles^63,64^. This design assessed prediction across observed feature–batch contexts, not extrapolation to wholly unseen proteins.

### Reference-anchored integration

Sample-to-reference ratios (SRRs) were calculated from Baseline-tier measurements on the log₂ scale as shown below, where the reference centre was the arithmetic mean of the finite reference measurements within each batch and feature. P was the primary anchor and N was evaluated as an alternative. Sample-level effects were assessed by PCA and SNR, feature-level reproducibility by two-way absolute-agreement single-measure intraclass correlation coefficients, and cross-platform effects by fold-change concordance. Global integration was assessed by sMAPE across non-reference samples. This ratio-based design followed the general common-reference principle established with Quartet materials^38^.

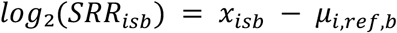

All 66 pairwise combinations of the 12 high-throughput analytical batches were evaluated under Balanced, Partial and Confounded sample-overlap designs. Balanced batches contained M, Y, P, X, F and N; Partial batches shared Y, P, X and N; Confounded batches separated M/Y from X/F while retaining P and N as potential bridges. Under Confounded designs, P-anchored methods were validated using N and N-anchored methods using P; Native and reference-free methods were evaluated in parallel P-and N-validation tracks. The 20 pipelines comprised Native; five reference-free batch-effect corrections (MAD scaling, quantile normalization, ComBat, RUV-III-C and RUVg); P-and N-informed RUVs; P-and N-anchored SRR; and ten SRR-plus-reference-free combinations^65–68^.

Eligible features required unambiguous protein mapping, finite M and F arithmetic means supported by at least two replicates in both batches, finite MAPD estimates and an M/F difference exceeding 3×MAPD in at least one batch. Complete measurements were required across design-available samples in both Baseline and final branches; no imputation was performed. Corrected values were evaluated on the linear scale. Quantitative agreement required sMAPE≤30%. Expected-response retention required finite Y and X TRCs, mean absolute deviation <0.25 and strict monotonicity of the pooled M–Y–X–F trajectory. A feature was harmonized only when both criteria were satisfied. Method-level rates were summarized by the median across batch pairs. Strategy success required at least one constituent method to harmonize the feature. Balanced-design consensus success was the proportion of 20 pipelines succeeding for each feature–batch-pair combination and was aggregated pair first; zero-success features were unresolved. Reference-choice and blank-subtracted SRR analyses used the same criteria.

Within each platform-pair scenario, high-consensus proteins were those at or above the 75th percentile of Balanced-design consensus success, whereas failed proteins had zero success; intermediate proteins were excluded. Native measurement ranks and Human Protein Atlas blood concentrations were compared descriptively. Physicochemical properties were compared by two-sided Wilcoxon rank-sum tests with Cliff’s delta, requiring at least five proteins per group; Benjamini–Hochberg-adjusted P values were calculated separately within each platform-pair scenario.

### Statistics and reproducibility

All statistical tests were two-sided. Technical replicates were repeated measurements of the same Plasmix material within an analytical batch and were not treated as independent biological replicates. Plasma donations were pooled before Plasmix construction; individual donors therefore did not constitute biological replicates in downstream platform comparisons.

Analyses were performed using R 4.6.1 and Bioconductor 3.23. Principal analysis packages included limma 3.68.4, impute 1.86.0, brms 2.23.0, irr 0.85, effsize 0.8.1, clue 0.3-68, metap 1.14, clusterProfiler 4.20.0 and org.Hs.eg.db 3.23.1. Batch-correction implementations used sva 3.60.0, RUVSeq 1.46.0, RUVIIIC 1.0.19 and limma 3.68.4. Predictive modelling used randomForest 4.7-1.2, xgboost 3.2.1.1, lightgbm 4.7.0 and pdp 0.8.3. Protein-property preparation used Peptides 2.4.6, bio3d 2.4.5 and FreeSASA 2.2.1 through Python 3.9.6. Data handling and visualization principally used data.table 1.18.4, tidyverse 2.0.0, ggplot2 4.0.3 and ComplexHeatmap 2.28.0. Package versions were exported directly from the analysis environment; the software-environment record and analysis code are archived with the study.

## Data availability

Processed protein abundance profiles, sample metadata, feature metadata and source data used for downstream analyses are available on Figshare at https://doi.org/10.6084/m9.figshare.32797509.

## Code availability

All custom code used for data processing, statistical analyses and figure generation is available at https://github.com/lyaqing/plasmix-proteomics.

## Acknowledgements

We thank all volunteers who donated blood for the development of the Plasmix reference materials. We acknowledge the computing resources provided by Computing for the Future at Fudan (CFFF) and the Human Phenome Data Center of Fudan University. We thank Dr. Lei Li of the Core Facility Center, Capital Medical University, for valuable assistance with LC–MS/MS analysis. Some illustrations were created with BioRender.

## Author contributions

Yuanting Zheng and L.S. designed the study. R.L. provided access to clinical facilities for blood collection, and W.H. provided facilities for on-site sample processing. H.W. and Q.W.C. coordinated participant recruitment and blood collection; H.W. and W.H. subsequently prepared and aliquoted the source plasma pools. Y.L., Q.C.C. and Y.Y.Z. completed the formulation, aliquoting, storage and distribution of the reference materials, with support from X.L., Y.G. and Z.A. S.Z. provided quality assurance. W.X., J.Y., W.Y., X.B.Y., J.W., Z.F., X.Y. and T.G. contributed to data generation and associated quality control. Q.C.C. and Yutong Zhang contributed to preliminary data analysis, review and verification. Y.L. developed the analytical methodology, performed the formal analyses, and prepared the figures and source data. Y.L., Yuanting Zheng and L.S. interpreted the results. W.X., Z.H.L., H.S., Z.Y., X.P.Y., G.L., Y.Y., Z.P., R.Z., J.M.L., X.F., L.J., C.D., F.H., Q.T. and X.C. contributed to discussions of the study. Y.L. wrote the original draft. Yuanting Zheng and L.S. revised the manuscript, supervised the study and acquired funding. All authors reviewed and approved the final manuscript.

## Extended data figure legends

**Extended Data Fig. 1.**
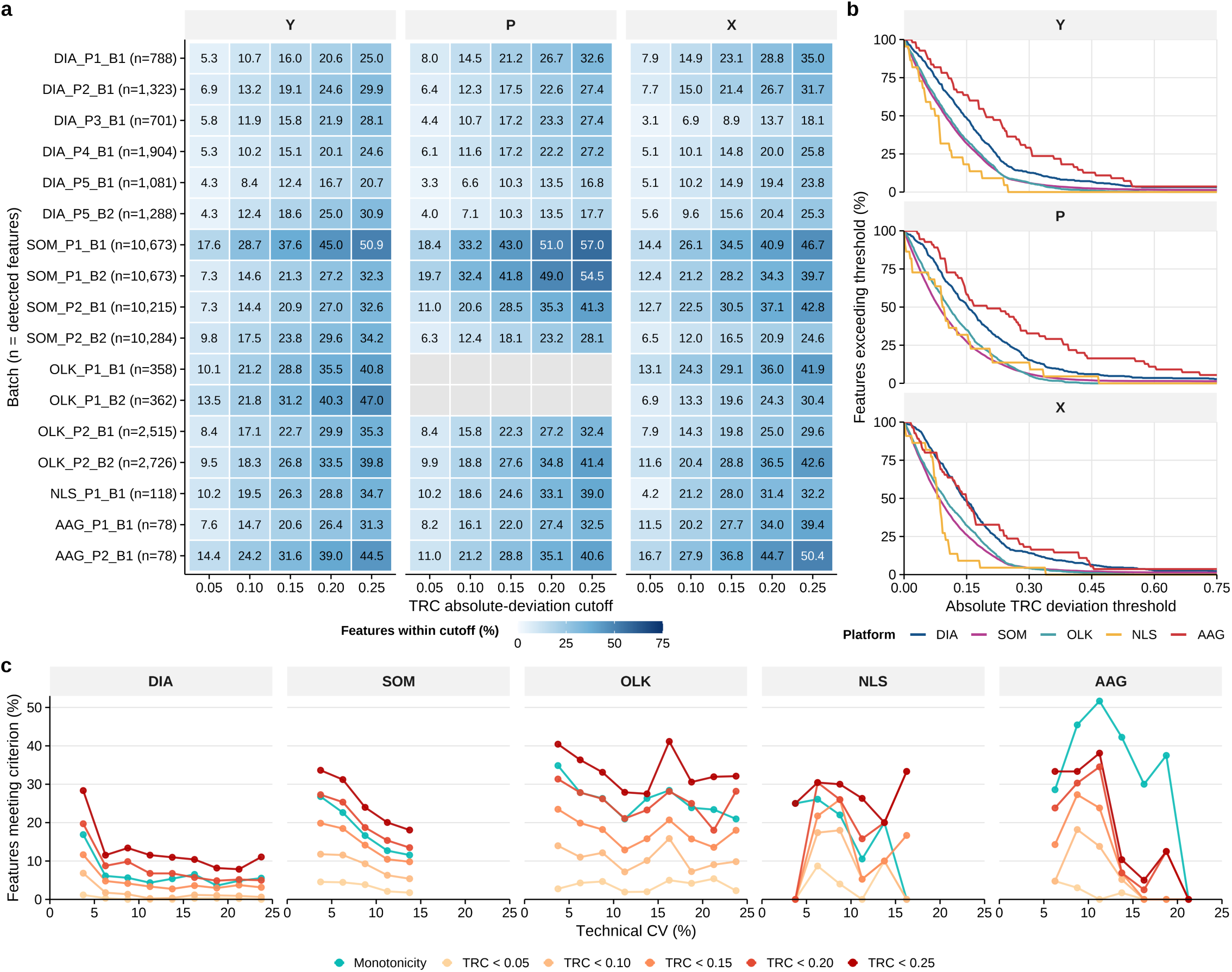
Titration-response criteria and technical precision. **a,** Gradient-specific sensitivity of absolute TRC deviation to the selected cutoff. For feature *i* and intermediate gradient sample 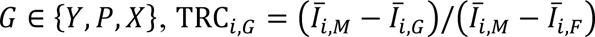, where *I* denotes the back-transformed group centre. Nominal TRC positions were 0.25, 0.50 and 0.75 for Y, P and X, respectively. Columns indicate absolute-deviation cutoffs from 0.05 to 0.25. Cell values show the mean percentage of detected features below each cutoff across valid three-replicate combinations. **b,** Complementary cumulative distributions of gradient-specific absolute TRC deviation among detected features satisfying titration monotonicity. At each threshold, curves show the percentage of feature–batch observations with an absolute deviation greater than or equal to that threshold. Panels correspond to the Y, P and X intermediate samples. **c,** Relationships between technical CV and titration-response criteria. Detected feature–batch observations were grouped into 2.5-percentage-point CV intervals. Curves show the batch-averaged percentages satisfying titration monotonicity or mean absolute TRC-deviation cutoffs from 0.05 to 0.25. Within each batch, intervals representing <2.5% of evaluable features were excluded, and platform-level points required support from at least half of the available batches. CV intervals below 25% are shown.

**Extended Data Fig. 2.**
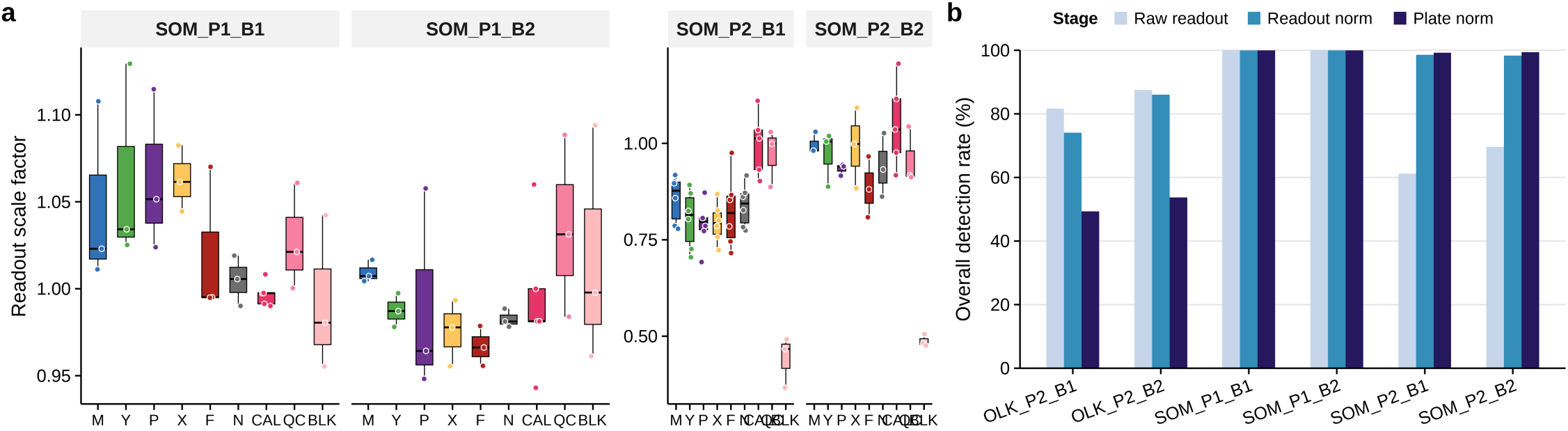
Stage-specific normalization and feature detection in affinity-based assays. **a,** Readout-normalization scale factors across M, Y, P, X, F, N, calibrator (CAL), quality-control (QC) and buffer-blank (BLK) samples in four SOMAmer-based batches. Boxes show medians and interquartile ranges; points denote individual samples. **b,** Overall feature-detection rates at the Raw readout, ReadoutNnorm and PlateNorm stages for two Olink and four SOMAmer-based batches. At the Raw readout and ReadoutNnorm stages, feature-and batch-specific thresholds were the BLK median plus three median absolute deviations on the linear scale. Plate norm used the platform-specific calibrated-tier detection status. A feature was counted as detected when more than half of the structurally available replicates passed in at least one of M, Y, P, X, F or N.

**Extended Data Fig. 3.**
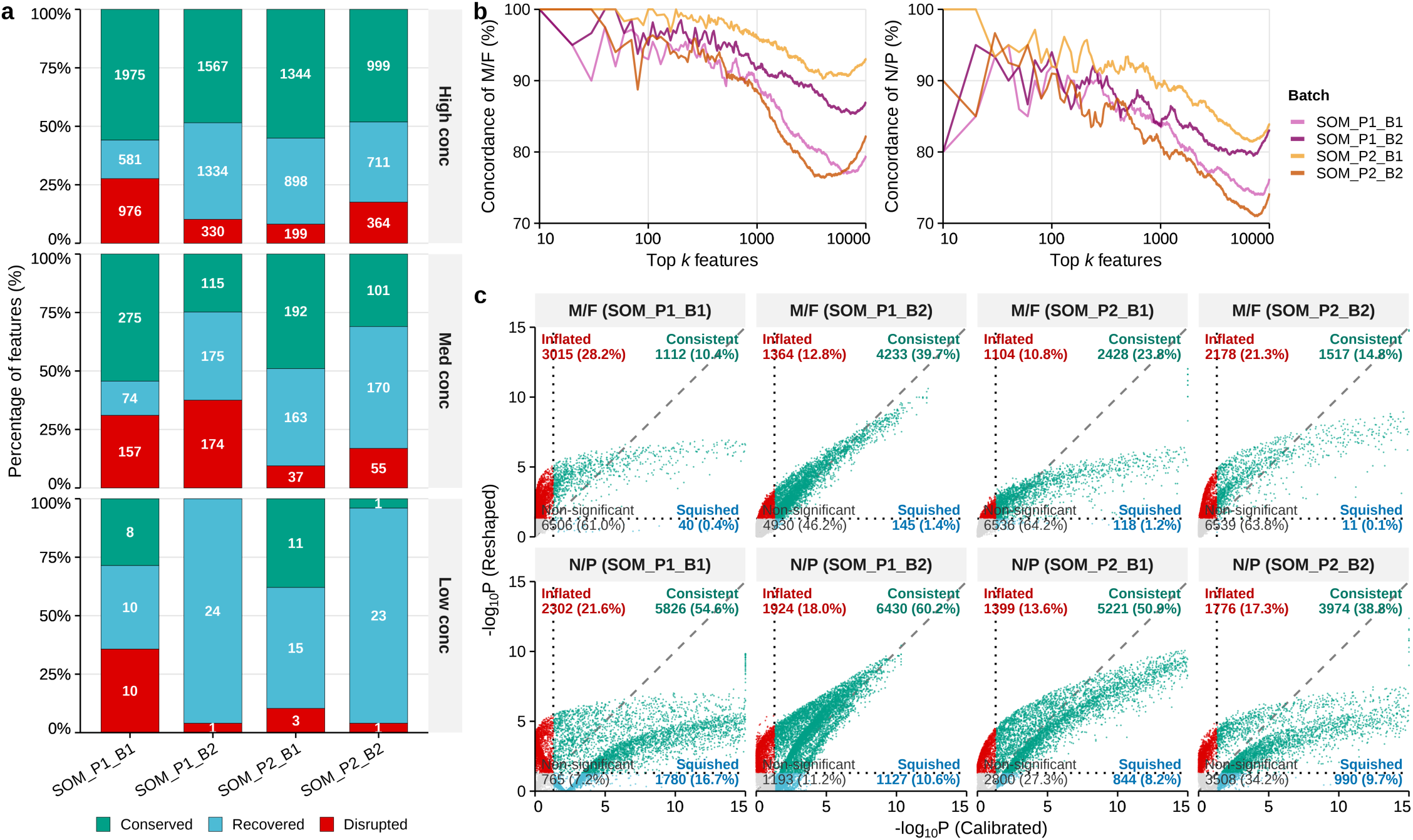
Adaptive normalization reshapes SomaScan titration responses. **a,** Expected-response transitions from calibrated to reshaped outputs, shown by SomaScan batch and SOMAmer dilution tier. The analysis includes features meeting the expected-response criteria at one or both stages. Conserved denotes features meeting the criteria at both stages; Recovered, only after reshaping; and Disrupted, only before reshaping. Numbers within segments denote feature counts. **b,** Concordance-at-the-top analysis of calibrated and reshaped outputs for the M/F and N/P contrasts. Features were ranked separately by absolute log₂ fold change. Curves show the percentage of the top *k*positions occupied by features present in both top-*k* sets and having the same fold-change direction. Colours denote batches. **c,** Comparison of −log₁₀-adjusted *P*values between calibrated and reshaped outputs. The dashed diagonal denotes equal values, and dotted lines denote adjusted *P* = 0.05. Points are classified as Consistent, significant at both stages; Inflated, significant only after reshaping; Squished, significant only before reshaping; or Non-significant, significant at neither stage. Numbers show feature counts and percentages within each panel.

**Extended Data Fig. 4.**
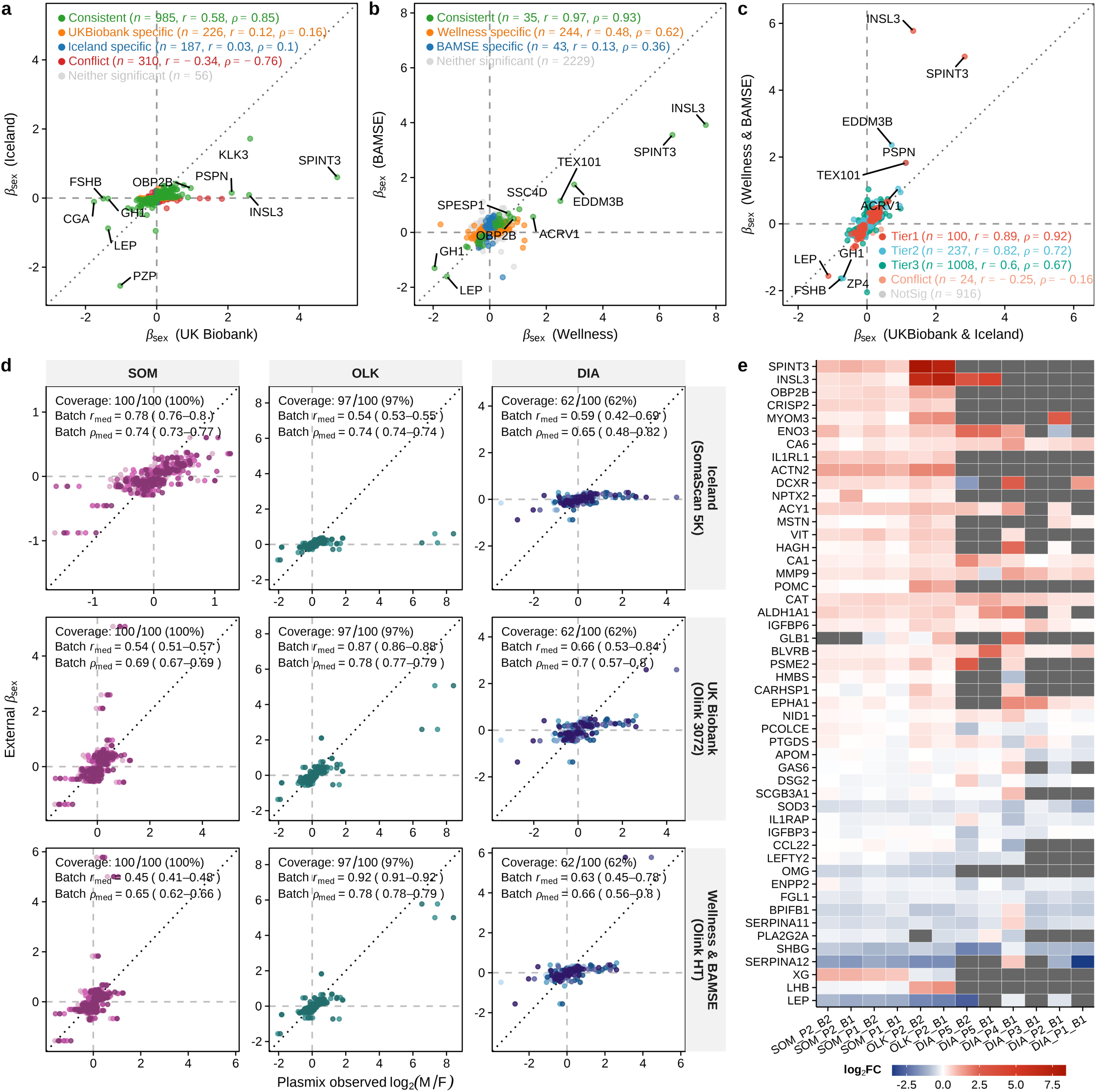
Consistency of sex-associated proteins across cohorts and Plasmix. **a,** Comparison of sex-associated effect estimates (*β*_sex_) between UK Biobank, measured using Olink Explore 3072, and Iceland, measured using SomaScan 5K. **b,** Comparison of *β*_sex_ between Wellness and BAMSE, both measured using Olink Explore HT. In **a,b**, colours distinguish concordant, conflicting, cohort-specific and non-significant effects; annotations give protein counts and Pearson *r*and Spearman *ρ*. **c,** Comparison of mean *β*_sex_ estimates from UK Biobank and Iceland with those from Wellness and BAMSE, stratified by evidence tier. **d,** External *β*_sex_ estimates versus Plasmix log₂(M/F) effects for Tier 1 proteins, stratified by external and Plasmix platforms and coloured by Plasmix batch. Annotations give Tier 1 coverage and batch-level correlation summaries. **e,** Plasmix log₂(M/F) effects across 12 batches for the 50 Tier 1 proteins with the largest absolute effects among proteins represented on at least two platforms. Cohort support required FDR < 0.05 and ∣ *β*_sex_ ∣≥ 0.05. Tier 1 required support in at least three cohorts and across Olink and SomaScan; lower tiers denote fewer cohort or platform supports, and Conflict denotes opposing supported effects. Positive values indicate higher abundance in males.

**Extended Data Fig. 5.**
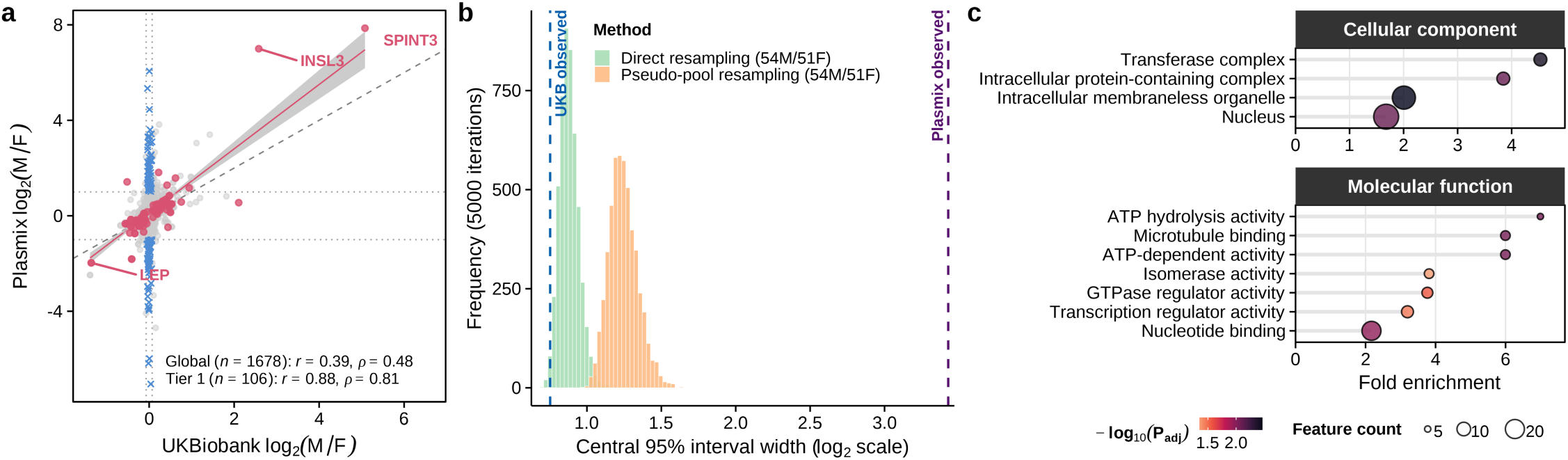
Magnitude and characteristics of the Plasmix sex gradient. **a,** UK Biobank versus Plasmix log₂(M/F) effects for matched Olink proteins measured using Olink Explore 3072 and Explore HT, respectively. Tier 1 and excess-shift proteins are highlighted; dotted lines mark excess-shift thresholds and the dashed line marks equality. The fitted line is shown with its 95% confidence interval, and annotations give Pearson *r*and Spearman *ρ*. Excess shift was defined as ∣ UKB log_2_(M/F) ∣≤ log_2_(1.05) and ∣ Plasmix log_2_(M/F) ∣≥ 1. **b,** Central 95% interval widths of UK Biobank log₂(M/F) effects across 5,000 resampling iterations using 54 male and 51 female participants and the same matched protein set as in **a**. Direct resampling used the difference between mean male and female NPX values, whereas pseudo-pool resampling averaged linearized NPX values within each sex before returning to the log₂ scale. Each width was defined as the 97.5th–2.5th percentile span of protein-level effects. Histograms show the resulting width distributions; dashed lines mark the observed UK Biobank and Plasmix widths. **c,** Gene Ontology enrichment of excess-shift proteins using all matched UK Biobank–Plasmix proteins as the background. Dot position, size and colour indicate fold enrichment, protein count and adjusted *P* value, respectively.

**Extended Data Fig. 6.**
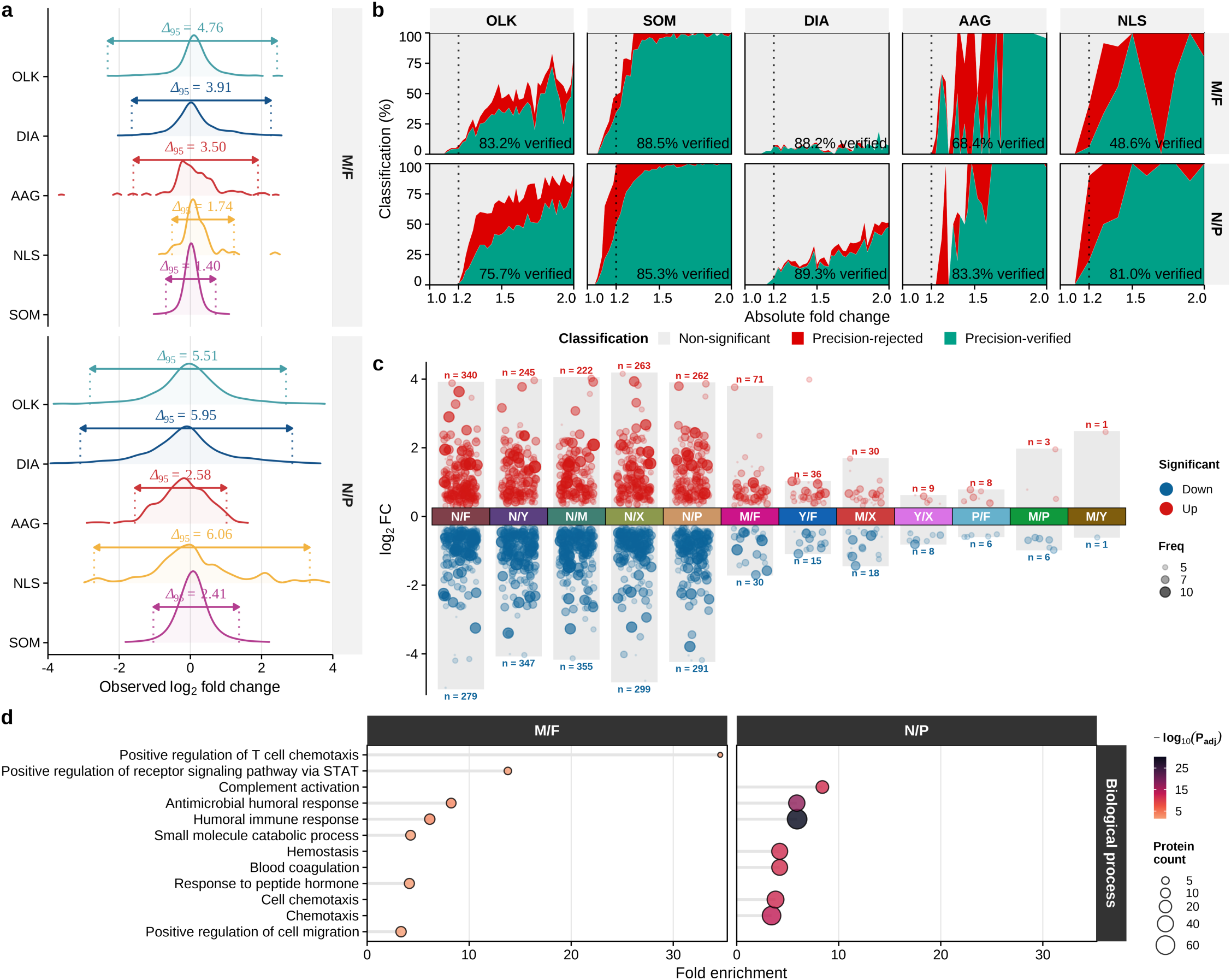
Platform-dependent fold changes across sample contrasts. **a,** Distributions of observed log₂ fold changes for the M/F and NIST–Plasmix N/P contrasts across five platforms. Annotations show the central 95% interval widths. **b,** Proportions of non-significant, precision-rejected and precision-verified observations across absolute fold-change bins. Precision verification required FDR < 0.05 and an absolute effect exceeding three times the feature-specific median absolute pairwise difference (MAPD). Dotted lines mark a 1.2-fold change; annotations give precision-verified proportions among significant observations above this threshold. **c,** Consensus differential effects across pairwise Plasmix and NIST contrasts. Consensus proteins were supported in the same direction by at least two platforms and four analytical observations. Points show median log₂ fold changes, with size and opacity denoting the number of supporting observations; labels give upregulated and downregulated protein counts. **d,** Gene Ontology biological-process enrichment of consensus M/F and N/P proteins, using all harmonized measured proteins with valid Entrez mappings as the shared background. Semantically related terms were simplified at a similarity cutoff of 0.65, retaining the term with the smallest adjusted *P* value within each similarity cluster. The top-ranked significant terms, ordered by adjusted *P* value, are shown for each contrast. Dot position, size and colour indicate fold enrichment, protein count and adjusted *P* value, respectively.

**Extended Data Fig. 7.**
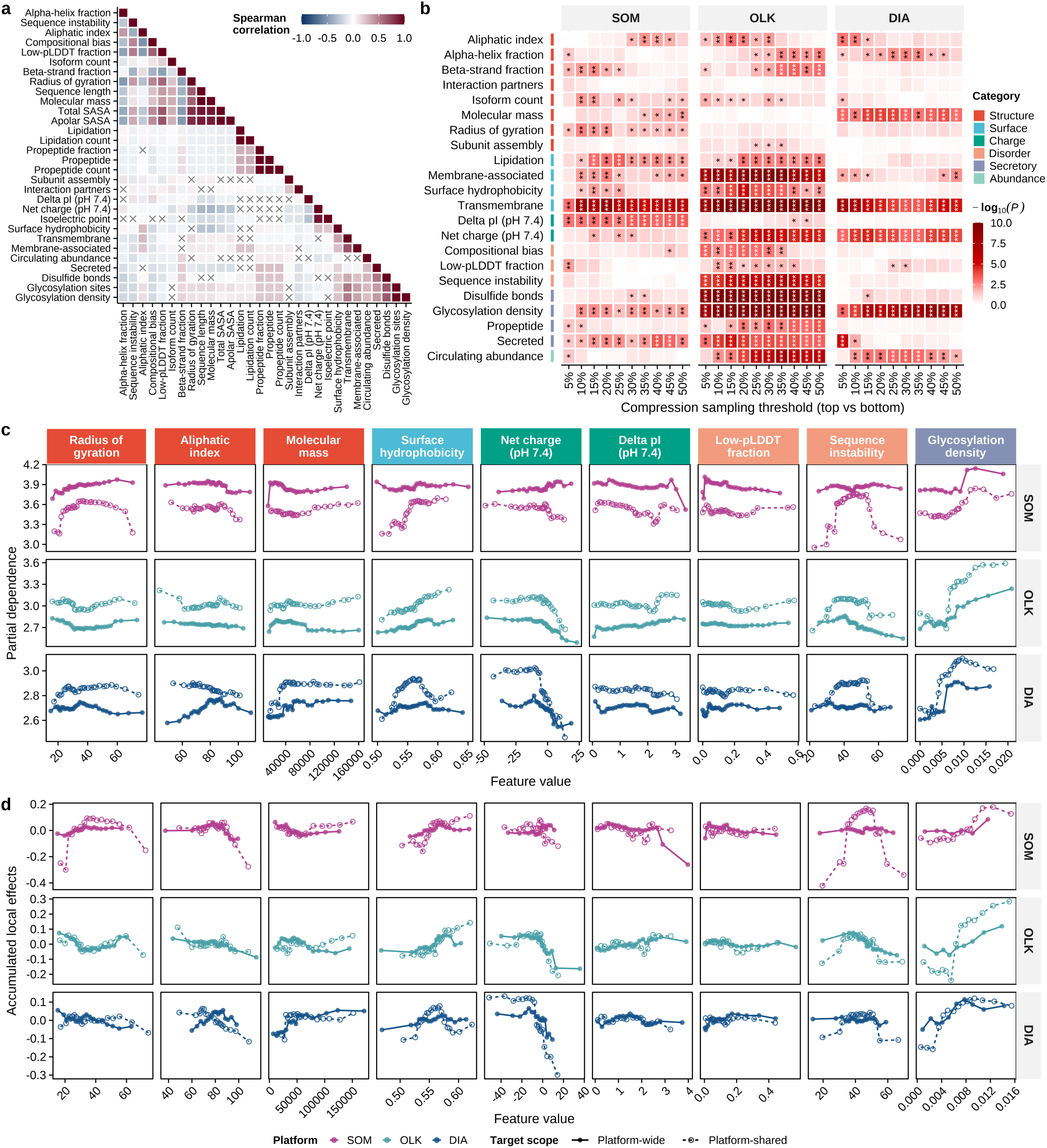
Physicochemical correlates and model-derived patterns of quantitative distortion. **a**, Spearman correlation matrix of physicochemical properties, ordered by Ward.D2 hierarchical clustering using 1−∣ *ρ* ∣. Crosses denote unadjusted *P* > 0.05. **b**, Associations between physicochemical properties and relative expansion among platform-wide targets. At thresholds from 5% to 50%, property distributions were compared between the upper and lower tails of relative expansion using two-sided Wilcoxon rank-sum tests. Fill denotes −log₁₀(*P*), capped at 10 for display; \*\**P* < 0.001, \**P* < 0.01 and *P* < 0.05. **c**, One-dimensional partial-dependence profiles for the properties displayed in Fig. 4g, derived from platform-wide and platform-shared Random Forest models. PDPs are shown on the original expansion scale. **d**, Accumulated-local-effect profiles for the same properties and models, shown on the log₂ scale used for model fitting. Colours denote platforms; solid lines and filled points denote platform-wide targets, and dashed lines and open points denote platform-shared targets.

**Extended Data Fig. 8.**
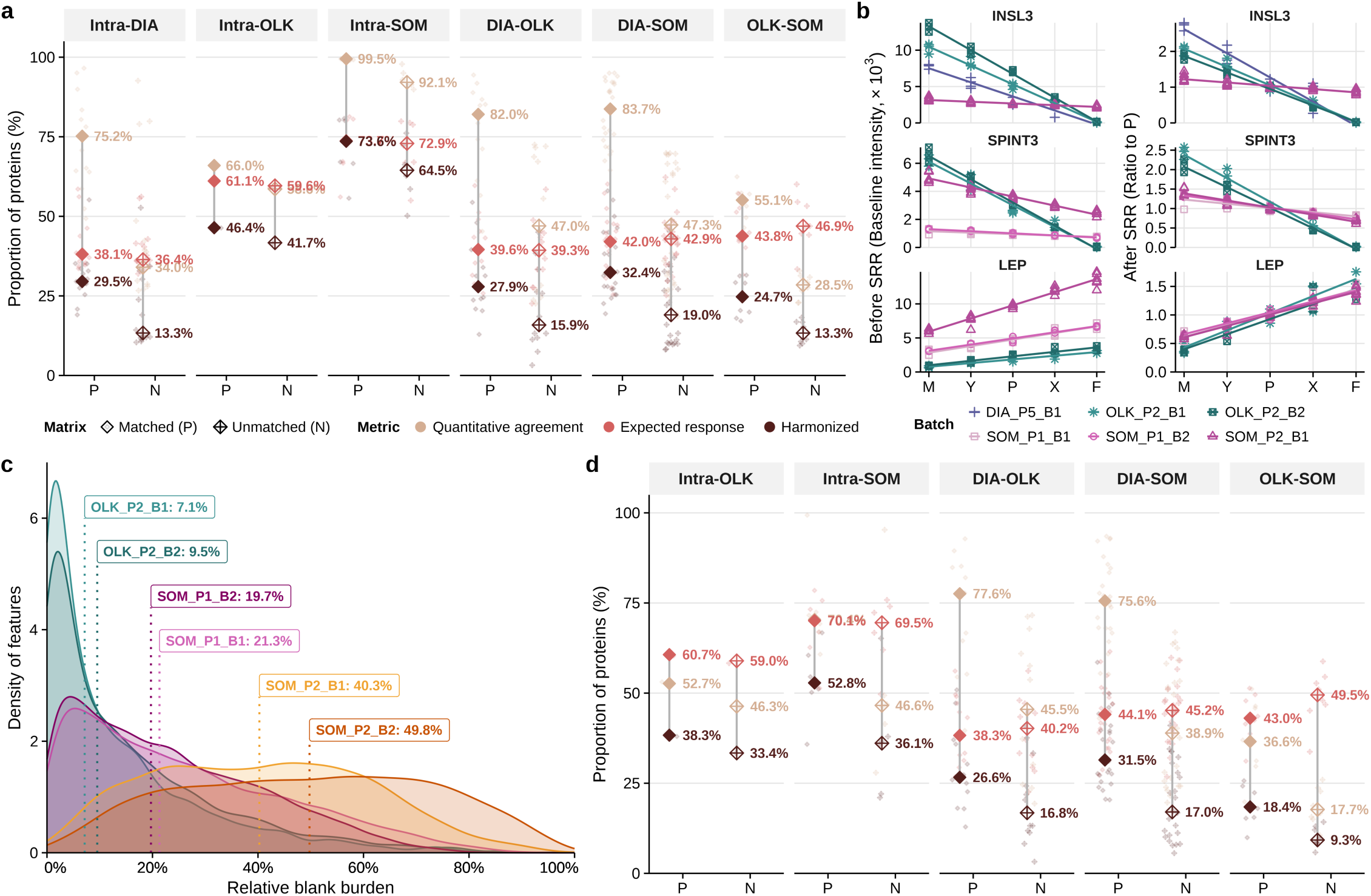
Reference choice, representative responses and blank subtraction in SRR integration. **a,** Balanced-design proportions of eligible proteins achieving quantitative agreement, expected response and harmonized status after standard SRR transformation using the matrix-matched Plasmix reference P or the unmatched NIST SRM 1950 reference N. Quantitative agreement required mean sMAPE ≤ 30% across M, Y, X and F; expected response required a monotonic M–Y–X–F trajectory and mean |TRC deviation| < 0.25; harmonized denotes joint passage of both criteria. Small points denote batch pairs and large symbols their medians. **b,** INSL3, SPINT3 and LEP trajectories across M, Y, P, X and F before and after SRR(P) transformation. Points denote technical replicates and lines denote batch-specific linear fits retained at R² > 0.6. Before-SRR values are linear Baseline-tier measurements; after-SRR values are ratios to the within-batch mean of P. **c,** Distributions of relative blank burden in Olink and SOMAmer-based batches, calculated for each feature as the median Raw-readout BLK measurement divided by the median Raw-readout P measurement. Only features detected in P at the Raw-readout stage were included. Dotted lines and labels indicate batch medians. **d,** Balanced-design quantitative-agreement, expected-response and harmonized rates after BLK-subtracted SRR(P) or SRR(N), using the same eligible feature sets as in a. Affinity-platform Raw-readout BLK measurements were subtracted before mapping back to the Baseline tier and calculation of sample-to-reference ratios. Intra-DIA comparisons were not evaluated because corresponding buffer-blank measurements were unavailable; DIA measurements were retained unchanged in cross-platform comparisons. Small points denote batch pairs and large symbols their medians.

**Extended Data Fig. 9.**
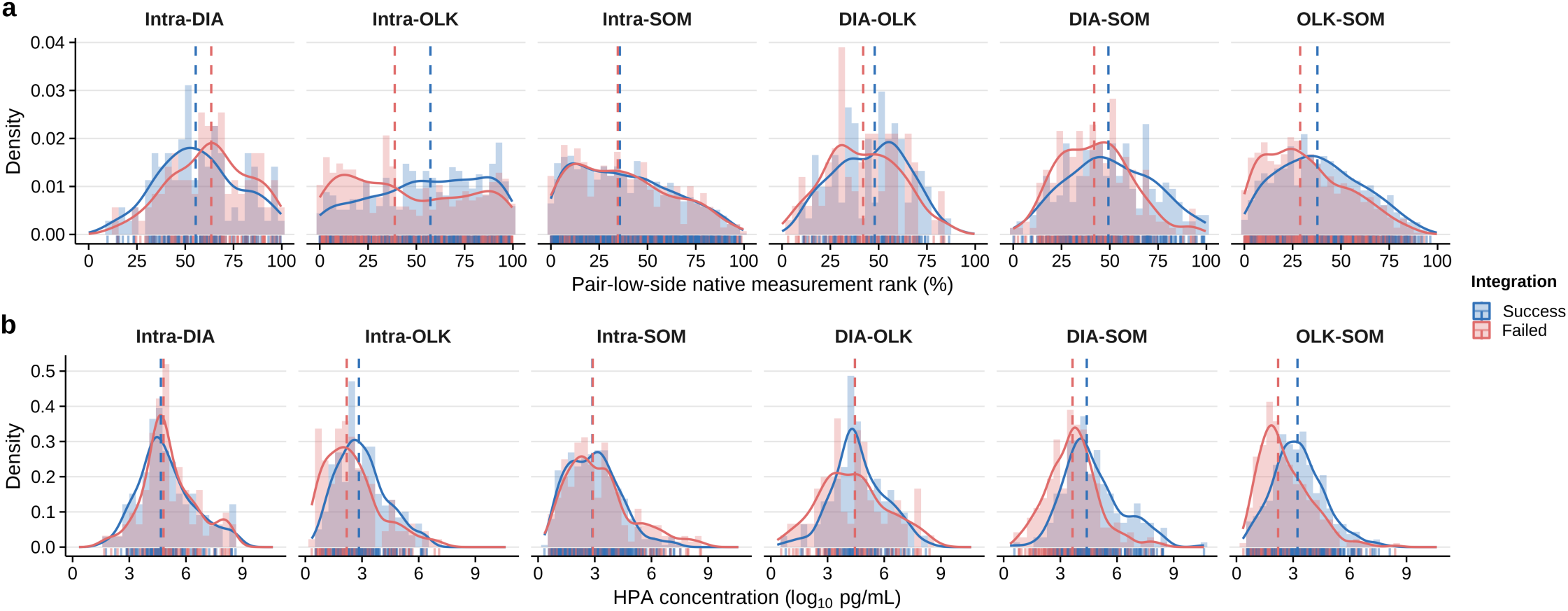
Native measurement rank and circulating abundance by integration outcome. **a,** Distributions of pair-low-side native measurement rank for upper-quartile consensus-success and zero-success proteins across the six integration scenarios. Within each batch pair, the lower of the two batch-specific measurement-rank percentiles was retained and then summarized by the median across evaluable pairs. **b,** Distributions of HPA-estimated blood concentration for the same protein groups, expressed as log₁₀ pg ml⁻¹. For each scenario under the Balanced design, consensus success was the median across batch pairs of the proportion of 20 pipelines satisfying both quantitative-agreement and expected-response criteria. Success denotes proteins at or above the scenario-specific 75th percentile, Failed denotes proteins with 0% consensus success, and intermediate proteins were excluded. Histograms and density curves show the distributions, rugs show individual proteins, and dashed lines mark group medians.

