## Supplementary Information for "Plasma titration provides a physical ruler for cross-platform proteomics"

Yaqing Liu et al.

### Reference materials

#### Plasmix reference materials

The study protocol was approved by the Ethics Committee of the School of Life Sciences at Fudan University (Approval No. FE253741). We recruited 110 healthy Chinese volunteers (55 males, 55 females; aged 20–60 years) with no known diseases and no medication use within the preceding week. Female participants were confirmed to be non-menstruating at the time of collection. Following an overnight fast, venous blood was collected into EDTA-$K_2$ vacuum tubes. Plasma separation was completed within 4 hours using a two-step centrifugation protocol: initial centrifugation at 3,000 × g for 15 min, followed by secondary centrifugation of the supernatant at 16,000 × g for 10 min.

Plasma donations were obtained from 108 of the 110 recruited participants and underwent quality assessment. Samples were screened for four blood-borne pathogens (HBsAg, HCV, TP, and HIV), and only negative samples were retained. Additionally, cell-free DNA was extracted from each donation for qPCR targeting the Y chromosome-specific SRY gene to verify the reported biological sex. One donation failed pathogen screening and two failed SRY-based sex verification; these three donations were excluded. The final qualified set comprised 105 donors (54 men and 51 women), yielding 1,790 mL and 1,690 mL of male and female plasma, respectively.

Qualified units were pooled by sex to create primary male (M) and female (F) pools. These base pools were volumetrically engineered to produce three intermediate mixtures with male-to-female ratios of 3:1 (sample Y), 1:1 (sample P), and 1:3 (sample X). All five reference materials (M, Y, P, X, F) were aliquoted and cryopreserved at –80 °C. The expected sex-specific expression gradient was independently validated via SRY qPCR quantification post-production.

#### NIST SRM 1950 reference material

We included the commercially available NIST Standard Reference Material® 1950 (Metabolites in Frozen Human Plasma), which is widely recognized as a metrologically traceable standard in metabolomics. This material consists of plasma pooled from 100 fasted, healthy donors (50 males and 50 females; aged 40–50 years) with an ethnic distribution representative of the U.S. population. Blood was collected into lithium heparin tubes, processed within 60 min of blood collection under controlled conditions, and subjected to centrifugation at 8,000 × g for 25 min before being aliquoted and stored at –80 °C.

### Platform-specific proteomic workflows

Quantitative protein profiles were generated across five distinct technology platforms encompassing affinity-based target recognition and mass spectrometry-based proteomics. Downstream data processing and relative abundance transformations were executed independently within each platform framework to preserve natively optimized parameters prior to multi-center integration. To eliminate position-based technical artifacts and prevent artificial sequential clustering, rigorous run-order strategies were strictly enforced. For the Olink, NULISA, and SomaScan assays, full-plate randomization was implemented across the assay plates following the assignment of fixed positions for manufacturer-specified reference calibrators, quality controls, and blank controls. For mass spectrometry workflows, samples were injected sequentially in structured loops encompassing the entire Plasmix suite (*i.e.*, continuous loops of M-Y-P-X-F-N), ensuring that technical replicates and biologically related samples were analyzed non-consecutively. For the AAgAtlas microarray, each sample was evaluated across three independent tubes, with each tube analyzed in consecutive technical triplicates within the array blocks. The loading sequence was structured in three successive loops encompassing all candidate samples, sequentially analyzing the triplicate blocks of the first, second, and third tubes respectively.

**Mass spectrometry-based proteomics**

Multiple pre-analytical enrichment protocols encompassing high-abundance protein depletion, neat plasma digestion, nanoparticle enrichment, and magnetic bead enrichment were evaluated across Orbitrap Exploris 480, Orbitrap Astral, and timsTOF Pro 2 hardware architectures operated in data-independent acquisition modes. Given the substantial biochemical baseline variations introduced by these divergent sample preparation mechanisms, raw files were intentionally not processed through a singular centralized pipeline. Instead, participating laboratories executed and delivered their natively optimized bioinformatics workflows, including the use of tailored homemade spectral libraries or in silico direct-DIA configurations, to maximize target protein coverage and quantification fidelity within their specialized analytical windows. Downstream multi-center integration was subsequently unified across the returned processed matrices. All granular batch-specific parameters, including chromatographic gradients, columns, flow rates, mass spectrometry source temperatures, hardware scan windows, and software configuration search parameters, are exhaustively cataloged in **Supplementary Table 17**.

**Olink proximity extension assay**

Multiplexed proteomic profiling was performed using the Olink Explore 384 Cardiometabolic and Explore HT panels. To accommodate the vast dynamic range of the plasma proteome, the two panels employ distinct physical processing strategies. While the Explore 384 architecture operates within a single analytical block, the Explore HT platform systematically fractionates each sample into five explicit dilution gradients directly mapped to eight physically segregated reaction blocks (1:1 for Blocks 1 through 4, 1:10 for Block 5, 1:100 for Block 6, 1:1,000 for Block 7, and 1:100,000 for Block 8). To rigorously monitor analytical quality and technical variance, three types of in-well internal controls (Incubation, Extension, and Amplification) were spiked into each sample on a per-block basis, while external controls consisting of three Negative Controls (NCs), three Plate Controls (PCs), and two Sample Controls (SCs) were distributed across fixed positions on every plate. Within this workflow, plasma samples were incubated with dual-recognition antibody pairs conjugated to unique DNA oligonucleotides. Upon proximal binding to the target proteins, these oligonucleotides hybridized and underwent polymerase-directed extension. The resulting amplicons were subsequently quantified via next-generation sequencing (NGS), employing an Illumina NovaSeq 6000 instrument for the 384 panel and an MGI DNBSEQ-T7 instrument for the HT panel. For data standardization, raw NGS counts for each target protein within individual samples were first normalized by dividing them by the counts of the sample-specific Extension Control uniquely assigned to its corresponding analytical block, and subsequently log2-transformed to generate intermediate ExtNPX values. Finally, inter-plate alignment was achieved by normalizing each sample's ExtNPX values against the plate-specific median of the pooled plasma Plate Controls on a strictly feature-by-feature basis, yielding the finalized relative Normalized Protein eXpression (NPX) metrics.

**SomaScan and Illumina Protein Prep assays**

High-plex aptamer-based profiling was performed using the SomaScan 11K Assay (v5.0) to capture approximately 11,000 protein measurements via Slow Off-rate Modified Aptamer (SOMAmer) reagents. To effectively capture the vast dynamic range of the plasma proteome, each plasma sample was physically fractionated into three distinct dilution groups (20%, 0.5%, and 0.005%) prior to assay incubation. Each standard 96-well assay plate accommodated 85 biological samples alongside 11 specialized plate controls, consisting of five Calibrators, three Quality Control (QC) samples, and three Buffer blanks. During the assay, 12 internal hybridization controls were incorporated into every sample well to account for readout efficiency. Following aptamer incubation and elution, target binding signals were captured via a fluorescence microarray-based readout hybridized to custom Agilent microarrays and scanned using a SureScan microarray scanner.

The downstream ADAT file standardization pipeline proceeded through sequential computational stages. Initial hybridization normalization (HybNorm) was applied independently to each sample using the specific in-well hybridization controls to correct for readout variances. This was followed by median intensity normalization within pooled calibrator replicates (MedNormInt), calculated independently for each dilution group. Subsequently, to eliminate inter-plate technical variance, the pipeline calculated the ratio of historical analyte-specific reference values to the intra-plate calibrator medians. This matrix of ratios was mathematically decomposed into two adjustment factors derived from the calibrators and sequentially applied as multipliers to all individual sample readouts on the plate: plate scaling (PlateScale), which computed a single median scaling factor per dilution group to remove bulk systematic intensity shifts, and plate-level calibration (Calibrate), which adjusted for remaining plate-specific variance on a strictly feature-by-feature basis utilizing analyte-specific calibration factors. Finally, Adaptive Normalization by Maximum Likelihood (ANML) was executed to normalize individual sample intensities by anchoring them to a pre-defined population reference. Assay quality strictly adhered to manufacturer criteria: all scale factors across the HybNorm, MedNormInt, PlateScale, and ANML stages were bounded within 0.4 to 2.5 thresholds. The Calibrate stage required the calibrator percent in-tails to be ≤ 10% outside the 0.6 to 1.4 range, while the QC check percent in-tails was mandated to be ≤ 15% outside the 0.8 to 1.2 range. ANML fraction utilization was required to be ≥ 30% within the 0.8 to 1.2 stabilization window.

The same proprietary SOMAmer chemistry was adapted to a high-throughput sequencing readout via the Illumina Protein Prep (IPP) 9.5K assay. Preserving the identical three-tier physical dilution strategy and 11-control plate layout configuration as the microarray baseline, the IPP workflow structurally diverged post-elution. To optimize sequencing efficiency and prevent hyper-abundant plasma proteins from monopolizing the sequencing flow cell, the protocol incorporated a proprietary Dynamic Range Compression (DRC) technology prior to library barcoding. This step utilized specific attenuator oligonucleotides (dummy probes) to compress read allocations from the highest-abundance analytes. The resulting libraries were sequenced on an Illumina NovaSeq X instrument, with demultiplexed reads processed through the DRAGEN Protein Quantification pipeline.

This computational framework replaced the population-scale ANML algorithm with a simpler median-based external normalization. Raw sequence counts were first normalized against internal spike-in controls to correct for sequencing readout variance, yielding the baseline ReadoutNorm values. The pipeline then streamlined data processing by consolidating the previous MedNormInt, PlateScale, and Calibrate steps into a unified external reference calibration stage (PlateNorm), followed by a final external median sample normalization (SampleNorm, historically referred to as MedNormExt). Technical quality control enforced a positive sample-level pass status and bounded all calculated scale factors (including HybNorm, PlateNorm, and SampleNorm) within the canonical 0.4 to 2.5 thresholds. At the PlateNorm stage, plates passed when <15% of SOMAmer scale factors fell outside the 0.6–1.4 range and when <15% of QC-check scale factors fell outside the 0.8–1.2 range.

**NULISAseq proximity ligation assay**

Targeted ultra-sensitive profiling was performed using the NULISAseq CNS Disease Panel 120 on the automated ARGO HT system. Each assay plate embedded four Negative Controls (NCs), three Inter-Plate Controls (IPCs), and three Sample Controls (SCs) alongside the biological samples. Within the ARGO HT system, the assay employed a dual-antibody liquid-phase reaction with intensive background-reduction washing, capturing target analytes via oligonucleotide-barcoded antibodies. Following the target capture reaction, an equal amount of exogenous mCherry internal control (IC) was spiked into every sample well to account for subsequent liquid handling and sequencing variances. The resulting barcoded libraries were subsequently quantified via next-generation sequencing employing an Element Biosciences AVITI system. Raw sequence FASTQ files were demultiplexed within the Alamar Control Center software. For data normalization, raw sequence counts were first adjusted against the specific in-well IC readouts, yielding IC-normalized counts. These values were subsequently normalized against the plate-specific IPCs. To calculate the final NULISA Protein Quantification (NPQ) metrics, these relative ratios were multiplied by 10,000, offset by adding a pseudo-count of 1 to accommodate zero-count readouts, and finally log2-transformed.

#### AAgAtlas autoantibody microarray

Distinct from the platforms quantifying circulating proteins, the AAgAtlas microarray was deployed to comprehensively profile the repertoire of circulating autoantibodies encompassing both IgA and IgG isoforms. Plasma samples were hybridized against thousands of full-length human proteins immobilized on the array surface, with autoantibody binding detected via Cy3-conjugated secondary antibodies for the IgG assay and Cy5-conjugated secondary antibodies for the IgA assay. One technical replicate of sample Y failed quality control and was excluded from subsequent analysis. Each sample was evaluated across three independent tubes, with each tube analyzed in consecutive technical triplicates within the array blocks. The loading sequence was structured in three successive loops encompassing all candidate samples, sequentially analyzing the triplicate blocks of the first, second, and third tubes respectively. Raw fluorescence images were scanned using GenePix Pro 7 software coupled with automated spot-localization software for automatic spot coordinates extraction. Quantitative signals were defined as the median fluorescence intensities at 532 nm and 635 nm (F532 median for the IgG assay and F635 median for the IgA assay, respectively) at each target coordinate. Data standardization was achieved by calculating the Signal-to-Noise Ratio (SNR) for each target, defined as the raw median spot intensity divided by a baseline background value derived from the 25th percentile of the signal intensity from all negative control spots on the respective array.

### Analytical feature annotation

#### Feature definition and protein mapping

To accommodate the distinct readouts across multiple technologies, we established a strict ontological framework separating platform-reported analytical proxies ("features") from biological entities ("target proteins"). Features denote the specific analytical identifiers (*e.g.*, individual SOMAmer probes, dual-antibody pairs, microarray antigens, or MS-derived protein groups) that index the final quantified measurements, distinguishing them from raw physical detector outputs (such as unassigned fluorescence emissions or MS precursor ions).

Based on their mapping to canonical UniProt accession numbers, features were classified into three categories. **Unique-target features (1:1 mapping)** exclusively target a specific protein, a stoichiometry strictly preserved across all Olink panels. **Co-targeting features (N:1 mapping)** occur when multiple distinct features independently target the same protein. This analytical multiplicity biologically reflects the detection of distinct structural epitopes (highly prevalent in SomaScan) or specific post-translational modifications, such as phosphorylation sites (observed in NULISA and AAgAtlas platforms). **Protein-group features (1:N mapping)** emerge when a single analytical feature maps to multiple distinct UniProt IDs. This ambiguity arises either from SOMAmers targeting multi-subunit protein complexes (annotated via hyphenated nomenclatures) or from MS workflows reporting indistinguishable protein groups (concatenated via semicolons) where specific proteoforms cannot be uniquely resolved.

To ensure unambiguous biological interpretation during downstream cross-platform integration, features lacking canonical UniProt IDs (unmapped) or classified as protein-group features (1:N) were strictly excluded. Only unique-target (1:1) and co-targeting (N:1) features were retained, yielding a final harmonized analytical space of 13,159 features.

#### Estimation of reference baseline concentrations

Analytical features were mapped to the Human Protein Atlas (HPA) blood reference database utilizing canonical UniProt accession numbers to estimate baseline absolute abundances. For target proteins possessing multiple database records, the median concentration value was extracted to mitigate individual outlier bias. The database cross-referencing prioritized quantitative mass spectrometry (MS) records; in instances where MS values were missing, immunoassay (IM) entries were utilized as the baseline reference. All retrieved values were uniformly converted to the clinical mass spectrometry consensus unit of pg/mL and log10-transformed. Features lacking unambiguous UniProt identifiers or corresponding HPA entries were excluded from abundance-dependent calculations.

#### Protein subcellular localization

Subcellular compartmentalization of target proteins was functionally annotated utilizing the HPA reference database. Classification was strictly rule-based: proteins featuring the term "membrane" in their class descriptor were classified as membrane proteins; those featuring "secreted" (exclusive of "membrane") were classified as secreted proteins. Targets possessing both annotations were categorized as secreted & membrane. Proteins lacking explicit annotation for either compartment were operationally classified as intracellular proteins for downstream analyses.

### Analytical data processing

**Hierarchical data tiers**

Given the diversity in proprietary data processing pipelines across technologies, a four-tier classification system was established to harmonize data structures for downstream analysis. **Tier 0 (Readout)** represents unprocessed detector outputs, including raw NGS read counts and initial fluorescence intensities.

**Tier 1 (Baseline)** comprises measurements corrected for basic technical noise or platform-specific internal-control effects. This tier encompasses DIA-MS LFQ intensities and AAgAtlas SNR, together with ExtNPX values for Olink, IC-normalized counts for NULISA, HybNorm values for SomaScan and ReadoutNorm values for IPP.

**Tier 2 (Calibrated)** encompasses data linearly transformed to achieve inter-plate standardization. This tier includes NPX for Olink, NPQ for NULISA, Calibrate data for SomaScan, and PlateNorm data for the IPP platform. The calibration mechanisms vary by technology: Olink and NULISA utilize internal controls (PCs and IPCs, respectively) to align readouts across plates, whereas the SomaScan and IPP pipelines compute feature-specific scale factors by evaluating plate-level calibrators against external historical reference datasets. Crucially, despite these methodological differences, all calibration adjustments within this tier are executed as feature-specific linear scalars applied uniformly across all samples within a plate. Consequently, Tier 2 strictly preserves the cross-sample relative quantitative rankings and original linear relationships for each individual analyte.

**Tier 3 (Reshaped)** is exclusively applicable to the SomaScan and IPP pipelines, specifically denoting their population-scale transformations (ANML and SampleNorm, respectively). These algorithms adjust individual sample distributions to match external reference populations. Unlike the purely feature-specific linear scaling utilized in Tier 2, these Tier 3 algorithms employ sample-by-sample adaptive scaling derived from external reference medians, thereby reshaping the native quantitative rankings among individual samples.

**Feature detection criteria**

Feature detection was evaluated at the Baseline tier for DIA-MS and AAgAtlas and at the Calibrated tier for Olink, NULISA and SomaScan. At the individual-measurement level, detection was defined using platform-specific criteria. For AAgAtlas, positive detection required an SNR $\geq 3$. For DIA-MS, detection required a non-zero, non-missing quantified intensity. For SomaScan 11K and IPP, the LoD in Buffer (LoDB) was calculated on the linear intensity scale as $\mathrm{median}(I_{\mathrm{BLK}}) + 3 \times \mathrm{MAD}(I_{\mathrm{BLK}})$, where $I_{\mathrm{BLK}}$ denotes the buffer-blank intensities and MAD denotes the median absolute deviation; the resulting LoDB was $\log_2$-transformed for comparison with the corresponding measurements. For NULISA, the LoD was calculated separately for each target on the normalized-count scale as $\mathrm{mean}(I_{\mathrm{NC}}) + 3 \times \mathrm{s.d.}(I_{\mathrm{NC}})$, where $I_{\mathrm{NC}}$ denotes the embedded negative-control measurements, and was subsequently converted to the NPQ scale using the same offset and $\log_2$ transformation applied to the sample measurements. For Olink, historical fixed background counts were mapped to the relative quantification space of the corresponding plate as $\log_2(\mathrm{LoDCount}/\mathrm{ExtCount}) - \mathrm{median}(I_{\mathrm{PC}})$, where LoDCount denotes the feature-specific fixed background count, ExtCount denotes the sample-specific Extension Control count and $I_{\mathrm{PC}}$ denotes the Plate Control intensities.

Within each analytical batch, a feature was considered detected in a sample group when more than half of its included technical replicates met the corresponding platform-specific criterion. A feature was classified as detected within the batch when this requirement was satisfied in at least one of the six study or reference sample groups (M, Y, P, X, F or N). Blank, calibrator and quality-control samples were not counted as analytical sample groups, although control measurements were used where required to define or map platform-specific detection thresholds.

For the titration-response, PCA-based SNR and technical-CV analyses in Fig. 2 and Extended Data Fig. 1, detection status was instead defined across the five Plasmix gradient samples (M, Y, P, X and F). NIST SRM 1950 was excluded from this definition because it was not part of the predefined titration series.

**Stage-specific detection thresholds**

Stage-specific feature detection was evaluated for Olink Explore HT and SOMAmer-based assays at the Raw readout, Readout norm and Plate norm stages. For Raw readout and Readout norm, feature- and batch-specific thresholds were calculated on the linear intensity scale as $\mathrm{median}(I_{\mathrm{BLK}}) + 3 \times \mathrm{MAD}(I_{\mathrm{BLK}})$, where $I_{\mathrm{BLK}}$ denotes the buffer-blank intensities; the resulting thresholds were $\log_2$-transformed for comparison with the corresponding measurements. For Plate norm, the platform-specific Calibrated-tier detection criteria described above were used. At each stage, a feature was considered detected in a sample group when more than half of its included technical replicates exceeded the corresponding threshold and was classified as detected overall when this criterion was met in at least one of M, Y, P, X, F or N. The overall detection rate for each batch and stage was calculated as the proportion of evaluated features classified as detected.

**Relative blank burden**

Relative blank burden (RBB) was evaluated in the Raw readout space for the same batches. For each feature and batch, RBB was calculated as $\mathrm{RBB} = \mathrm{median}(I_{\mathrm{BLK}})/\mathrm{median}(I_{\mathrm{P}})$, where $I_{\mathrm{BLK}}$ and $I_{\mathrm{P}}$ denote the linear Raw readout intensities of the buffer blanks and sample P, respectively. Only features for which sample P met the Raw readout detection criterion were retained. Batch-level RBB distributions and medians were used to summarize the contribution of blank background relative to the sample P signal.

**Missing value imputation**

We implemented a hierarchical imputation framework to maintain quantitative integrity across distinct analytical contexts. For differential expression analysis, missing values were imputed separately within each analytical batch, data tier and processing stage using a batch-specific low-intensity horizon. The horizon was defined as the 0.1th percentile of all finite values in the corresponding analysis matrix. Missing values were sampled from a Gaussian distribution centred on this horizon with an s.d. equal to 5% of its absolute value. For matrices containing only non-negative values, imputed values were bounded below at 50% of the batch minimum; for matrices containing negative $\log_2$-scale values, the lower bound was set to three s.d. below the horizon. Detection status and feature eligibility were determined from the original, unimputed measurements.

For global integration analyses, missing values were imputed using the k-nearest neighbours algorithm (KNN; k = 10). Before imputation, features were filtered according to cross-batch detection frequency. General global integration retained features detected in >50% of the evaluated batches. Global PCA retained features detected in >80% of the batches and detected in reference sample (P). KNN imputation was then applied to the remaining missing values. For PVCA, features missing across an entire batch were first imputed batch-wise using the LoD procedure. Remaining sporadic missing values were subsequently imputed using global KNN, yielding complete matrices for variance decomposition.

To facilitate principal variance component analysis (PVCA), we adopted a hybrid imputation strategy to address the dual mechanisms of missingness. Batch-wise LoD imputation was applied exclusively to features exhibiting complete batch missingness (Missing Not At Random, MNAR) to preserve platform-specific detection limits. Subsequently, the remaining scattered missing values (Missing At Random, MAR) were resolved using the global KNN algorithm. This two-stage workflow generates the complete data matrices required for variance decomposition while preserving the technical and biological variance signatures intrinsic to each batch.

**Differential expression analysis**

Differential expression analysis was conducted using limma separately for each analytical batch across the Baseline, Calibrated and Reshaped tiers and their available processing stages. Exactly three technical replicates were selected once for each eligible batch–sample group among M, Y, P, X, F and N using a fixed random seed and were retained consistently across processing stages and contrasts. For each contrast, features detected in at least one of the two compared sample groups were retained. Missing values were imputed using the low-intensity-horizon strategy described above, moderated test statistics were obtained using empirical Bayes methods, and P values were adjusted using the Benjamini–Hochberg procedure. Classification summaries across these tiers and processing stages are provided in Supplementary Table 6.

Extended Data Fig. 5a,b used the Baseline tier for DIA-MS and AAgAtlas and the Calibrated tier for Olink, SomaScan and NULISA; the consensus and enrichment analyses in Extended Data Fig. 5c,d used the same tier assignments but excluded AAgAtlas, as described below.

Feature-specific technical variation was estimated on the $\log_2$ scale separately within each analytical batch, data tier and processing stage. For each feature, the median absolute pairwise difference (MAPD) was calculated as the median of all absolute pairwise differences among technical replicates within the M, Y, P, X, F and N sample groups. Only sample groups with at least two available technical replicates contributed to MAPD estimation. Manufacturer-provided blanks, calibrators and quality-control samples were excluded because they did not represent the study or reference materials evaluated in the differential-expression analysis.

Feature-level results were assigned to four mutually exclusive categories: Non-significant (non-finite $\log_2$ fold change, adjusted P value or MAPD, or adjusted P value $\geq 0.05$); Precision-rejected (adjusted P value $< 0.05$ and $|\log_2$ fold change$| \leq 3 \times$ MAPD); Small-magnitude (adjusted P value $< 0.05$, $|\log_2$ fold change$| > 3 \times$ MAPD and $|\log_2$ fold change$| \leq \log_2 1.2$); and Verified-DEP (adjusted P value $< 0.05$, $|\log_2$ fold change$| > 3 \times$ MAPD and $|\log_2$ fold change$| > \log_2 1.2$). For visualization in Extended Data Fig. 5b, Small-magnitude and Verified-DEP results were combined into the Precision-verified category because both passed the feature-specific precision criterion.

### Titration-performance benchmarking

#### Analysis sets and replicate standardization

Primary titration-response analyses used Baseline-tier AAgAtlas SNR and DIA-MS intensities and Calibrated-tier Olink NPX, SomaScan Calibrate and NULISA NPQ outputs. SomaScan Reshaped-tier outputs were additionally retained for processing-stage comparisons. Detection status was treated as a separate annotation and did not determine whether a feature was numerically evaluable.

To standardize analyses across batches with different replicate depths while preserving acquisition-round structure, all distinct combinations of three acquisition rounds shared across eligible sample groups were enumerated. A sample group was eligible when at least three structurally available replicates were present; the M and F endpoint samples were additionally required for TRC and gradient-fit analyses. When an entire well was structurally unavailable according to the sample metadata, it was replaced by the nearest unused valid acquisition round. Feature-level missingness was not used to alter the replicate plan. The same replicate plans were used for the PCA-based SNR and technical-CV analyses, using the eligible Plasmix gradient groups available within each batch.

#### Performance assessment metrics

***Feature-level titration response coefficient.*** Within each three-replicate combination, the group-level $\log_2$ measurement, $\bar{L}_{i,s}$, was calculated as the mean of the finite measurements for feature $i$in sample group $s$. A group centre was considered evaluable when at least two of the three selected measurements were finite. No missing values were imputed for the feature-level TRC or monotonicity analyses. Group centres were converted to the linear scale as $I_{i,s} = 2^{\bar{L}_{i,s}}$. For each intermediate sample $s \in \{Y, P, X\}$, the titration response coefficient was defined as

$$\mathrm{TRC}_{i,s} = \frac{I_{i,M} - I_{i,s}}{I_{i,M} - I_{i,F}},$$

with nominal positions of 0.25, 0.50 and 0.75 for Y, P and X, respectively. The M and F centres were required to be finite and unequal. For each intermediate sample, TRC and absolute deviation from its nominal position were first averaged over all replicate combinations in which they were evaluable. The mean absolute TRC deviation for a feature was then calculated as the equally weighted mean across the available intermediate samples. At least two finite intermediate TRCs were required, and the TRC-deviation criterion was satisfied when the mean absolute deviation was $< 0.25$.

***Titration monotonicity and expected response.*** Monotonicity was evaluated using the TRC coordinate system, in which M and F were fixed at 0 and 1, respectively. For each feature, the sample groups with evaluable TRCs were ordered according to their nominal positions, and adjacent relations were defined dynamically from the available groups rather than requiring a complete five-sample series. Within each replicate combination, an adjacent relation was supported when the TRC of the earlier sample was strictly smaller than that of the later sample; ties were treated as failures. The support rate of each relation was calculated across replicate combinations in which both members of the relation were evaluable. A feature satisfied titration monotonicity when at least two adjacent relations were defined, every defined relation was evaluable in at least one replicate combination and every relation had a support rate $> 0.5$. A feature was classified as showing the expected titration response when it satisfied both the monotonicity criterion and the mean absolute TRC-deviation criterion described above. Detection status was not part of this classification.

***Batch-level gradient fit.*** Recovery of the expected mixture response was also assessed across features within each analytical batch and separately for Y, P and X. For feature $i$and intermediate sample $s$, the observed $\log_2$ response relative to F was

$$O_{i,s} = \bar{L}_{i,s} - \bar{L}_{i,F}.$$

Given the empirical endpoint difference

$$\Delta_i = \bar{L}_{i,M} - \bar{L}_{i,F},$$

the expected response under linear mixing on the abundance scale was

$$E_{i,s} = \log_2[\pi_s 2^{\Delta_i} + (1 - \pi_s)],$$

where the male fractions $\pi_s$ were 0.75, 0.50 and 0.25 for Y, P and X, respectively. Gradient-fit $R^2$ was calculated without fitting an additional slope or intercept:

$$R_s^2 = 1 - \frac{\sum_i (O_{i,s} - E_{i,s})^2}{\sum_i (O_{i,s} - \bar{O}_s)^2}.$$

At least three features with finite observed and expected responses were required. The calculation was performed within each replicate combination and summarized as the mean and s.d. across combinations. Values were generated for both all evaluable and detected features; Fig. 2 displays the detected-feature results.

***Signal-to-noise ratio.*** As defined in the previous Quartet study, PCA-based SNR quantified separation among the available Plasmix gradient groups relative to dispersion among technical replicates. For SNR only, pre-existing missing values were imputed at a batch-specific low-intensity horizon using the procedure described above; finite measurements below the platform-specific detection threshold were retained unchanged. Features with zero variance in the selected replicate combination were excluded. PCA was centred for all platforms, with unit-variance scaling applied to DIA-MS and no unit-variance scaling applied to SomaScan, Olink, NULISA or AAgAtlas data.

For samples $a$ and $b$, their distance in the first two principal components was calculated as

$$d_{a,b} = \sum_{k=1}^{2} w_k \left(z_{a,k} - z_{b,k}\right)^2,$$

where $z_{a,k}$ is the score of sample $a$ on principal component $k$, and $w_k$ is the proportion of variance explained by that component. SNR was then defined as

$$\mathrm{SNR} = 10\log_{10}\left(\frac{\mathrm{mean}\left(d_{a,b} \mid g_a \neq g_b\right)}{\mathrm{mean}\left(d_{a,b} \mid g_a = g_b\right)}\right).$$

SNR was calculated for every valid three-replicate combination and summarized as the mean and s.d. across combinations. The SNR used here is distinct from the AAgAtlas SNR representation derived from spot-to-background normalization.

***Coefficient of variation.*** Technical CV was calculated on the linear measurement scale without missing-value imputation. Within each three-replicate combination and sample group, all three selected feature measurements were required to be finite, and CV was calculated as the standard deviation divided by the mean. Feature-level CV within a replicate combination was obtained by averaging across the available Plasmix gradient groups, and the final feature-level CV was the mean across all valid replicate combinations. Fig. 2 summarizes the distributions of these feature-level CVs among detected features. AAgAtlas and selected Olink outputs were additionally displayed without detection filtering, and SomaScan Reshaped-tier values were included as processing-stage comparisons.

***Symmetric mean absolute percentage error (sMAPE).*** Cross-batch quantitative agreement was measured as the sMAPE between the two batches' arithmetic mean values for each evaluation sample, averaged across the design-specific shared or validation samples. Per-batch arithmetic means provided one group centre for each batch and sample. Because sMAPE for two batch means is proportional to their coefficient of variation by a factor of approximately $\sqrt{2}$, a 30% sMAPE threshold corresponds to an inter-batch CV of ~20%. This aligns with the lower limit of quantification (LLOQ) acceptance criteria defined by the FDA and ICH M10 bioanalytical guidances. The expanded 30% bound relative to a standard 15% intra-batch CV reflects the mathematical propagation of two independent per-platform tolerances across batches.

**Titration-response summaries and sensitivity analyses**

Titration-response performance was summarized at the feature–batch level using effect-size-stratified and rank-based analyses. Observations from the primary analysis tiers with finite M/F effects and defined expected-response classifications were grouped by platform and signed $\log_2$(M/F) intervals of width 0.2; intervals containing fewer than two observations were excluded. Expected-response rates were calculated separately for all evaluable and detected observations, and LOESS curves were fitted using a span of 0.8 and a second-degree local polynomial. In parallel, observations were ranked within each batch by descending absolute $\log_2$(M/F), and the cumulative expected-response rate was calculated at each rank for the all-evaluable and detected feature sets. Batch trajectories were extended to the maximum platform-specific rank by carrying forward their terminal values and were summarized across batches using the mean and, for detected features, the interquartile range.

Detection-adjusted recovery yield was defined to integrate analytical detection and quantitative-response performance without conditioning expected-response classification on detection. Within each batch, the denominator comprised the union of all detected features and non-detected features satisfying the expected-response criteria, whereas the numerator comprised the expected-response features within this set. Contributions from detected and LoD-excluded expected-response features were quantified separately. The same calculation was applied to SomaScan Reshaped-tier measurements for processing-stage comparisons.

Sensitivity analyses were restricted to detected features. Intermediate-specific TRC performance was evaluated across absolute-deviation cutoffs of 0.05, 0.10, 0.15, 0.20 and 0.25 and summarized across valid replicate combinations. Complementary cumulative distributions were calculated for absolute TRC deviations among features satisfying the titration-monotonicity criterion, with values exceeding 0.75 retained in the denominator. The relationship between technical CV and titration performance was evaluated among features with at least two finite intermediate TRCs, at least three completely evaluable adjacent relations, and finite CV and mean TRC deviation. CV values were grouped into 2.5-percentage-point intervals, and monotonicity and TRC-deviation rates were calculated within each batch and interval. Sparse batch-level intervals representing less than 2.5% of eligible features were excluded, and platform-level values were calculated as unweighted means across batches and retained when represented in at least half of the platform's batches.

**Correspondence-at-the-top analysis**

Correspondence-at-the-top (CAT) analysis was performed for the M/F and N/P contrasts across the 12 high-throughput batches using Baseline-tier DIA intensities, Calibrated-tier Olink NPX values and Calibrated-tier SomaScan values. $\text{Log}_2$ fold changes were obtained from the batch-level limma analyses. Features were mapped to canonical UniProt accessions, and proteins represented by more than one retained analytical feature within a batch were excluded from the corresponding protein-level comparison.

For each batch pair and contrast, the analysis was restricted to proteins quantified in both batches. Proteins were ranked separately in each batch by absolute $\log_2$ fold change. CAT correspondence was evaluated at top-ranked fractions from 1% to 100% of the common-protein universe, in increments of one percentage point. At each fraction, the corresponding number of proteins was selected from each ranked list, and correspondence was defined as the number of proteins simultaneously present in both top-ranked sets and showing the same effect direction, divided by the number selected from each list. Curves were averaged across a fixed set of eligible batch pairs within each comparison category, so that the contributing batch-pair composition remained unchanged across ranking fractions. Batch pairs were categorized as within protocol, across protocols within the same platform or across platforms.

Random-ranking expectations were calculated analytically and separately for each batch pair, contrast and ranking fraction using the actual common-protein universe and the observed proportions of positive, negative and zero effects in the two batches. For ranking fraction $f$, the expected correspondence was:

$$f \sum_{s \in \{-1,0,1\}} p_{1s}\, p_{2s}$$

where $p_{1s}$ and $p_{2s}$ denote the proportions of proteins with direction $s$in the two batches. Pair-specific expectations were subsequently averaged across the fixed batch-pair set contributing to each comparison category.

**Cross-setting profile concordance by expected-response status**

Cross-setting comparisons used the same 12 high-throughput batches and primary data layers as the expected-response analysis. The protein-level analysis included platform-detected features and features below the platform-defined detection threshold that nevertheless met the expected-response criteria. Finite M and F group centres were required. When multiple analytical features mapped to the same UniProt accession within a batch, that protein–batch record was excluded rather than averaged. For each batch pair, proteins were classified as joint pass, one pass or neither pass according to whether they met the expected-response criteria in both, one or neither batch.

Profile agreement was compared between joint-pass and neither-pass proteins. Within each batch, M and each commonly available intermediate sample were expressed as linear fold changes relative to F; F therefore had a fixed value of one and was not included in the error average. M and at least two intermediate samples shared by both batches were required. Profile sMAPE across M and the shared intermediate samples was calculated as:

$$\text{sMAPE} = 100 \times \text{mean}\left(\frac{|\, a - b \,|}{(|\, a \,| + |\, b \,|)/2}\right)$$

To control for differences in contrast magnitude, joint-pass and neither-pass proteins were optimally matched without replacement within each batch pair using the Hungarian algorithm. Matching minimized the absolute distance in $\log_1$p-transformed mean M/F effect magnitude, where the matching variable was the mean of the two absolute batch-specific $\log_2$(M/F) effects. No caliper or random subsampling was applied, and the larger group was reduced to the size of the smaller group. The panel reports, for each batch pair, the difference between the median sMAPE of matched neither-pass proteins and that of matched joint-pass proteins; positive values indicate lower profile error among joint-pass proteins.

### Biological validation

#### External validation in the China Kadoorie Biobank

Participant-level Olink–SomaScan correlations and protein–phenotype effect estimates were obtained from the published China Kadoorie Biobank supplementary data. Only strict one-to-one mappings among canonical UniProt accessions, Olink assay identifiers and SomaScan assay identifiers were retained. The Plasmix classification used two Olink Explore HT NPX batches and two non-ANML SomaScan Calibrate batches. Proteins were required to be represented in all four batches. The number of jointly passing Olink–SomaScan batch combinations was calculated as the product of the numbers of passing Olink and SomaScan batches. Supported proteins had at least two jointly passing combinations, whereas Absent proteins had none; proteins with one jointly passing combination were not included in the comparison.

Supported and Absent proteins were matched 1:1 without replacement by Hungarian optimization of $\log_1$p-transformed Plasmix M/F effect magnitude. Effect magnitude was calculated as the mean of the absolute median Olink and SomaScan M/F effects across their respective two batches. Matching used no caliper or random sampling. For participant-level agreement, the published Spearman correlation between Olink and non-ANML SomaScan measurements was extracted for each protein. The median paired difference between matched Supported and Absent proteins was summarized with a percentile 95% confidence interval from 1,000 bootstrap resamples.

For phenotype-effect concordance, the same matched protein sets were evaluated across 18 non-sex traits. Pearson and Spearman correlations between Olink and non-ANML SomaScan effect estimates were calculated separately for each trait using proteins with finite estimates on both platforms. Pearson correlations are displayed in Fig. 3e, and the corresponding Spearman results are provided in the Source Data.

**Consensus differential proteins and functional enrichment**

Verified-DEPs were integrated across analytical observations to identify cross-platform consensus differential proteins. The analysis included Baseline-tier DIA-MS data and Calibrated-tier Olink, SomaScan and NULISA data. AAgAtlas was excluded because it quantified autoantibody binding rather than circulating protein abundance. For each protein and contrast, a candidate direction was retained when it was supported by at least two technology platforms and four analytical observations. The integrated effect size was calculated as the median $\log_2$ fold change across supporting observations. *P* values were combined using Fisher's method implemented in the metap package and were subsequently adjusted across all candidate protein–contrast combinations using the Benjamini–Hochberg procedure. Consensus proteins were required to have an adjusted *P* value < 0.05 and |median $\log_2$ fold change| > $\log_2(1.2)$.

Gene Ontology over-representation analysis was performed for the Biological Process, Cellular Component and Molecular Function domains using clusterProfiler and org.Hs.eg.db. UniProt accessions were mapped to human Entrez identifiers, and all harmonized measured proteins with valid UniProt-to-Entrez mappings were used as the shared background universe for the M/F and N/P contrasts. Enrichment was evaluated using the compareCluster function, with Benjamini–Hochberg-adjusted P < 0.05 considered significant. Semantically related terms were simplified using the simplify function with a similarity cutoff of 0.65, retaining the term with the smallest adjusted P value within each similarity cluster. The top-ranked Biological Process terms were selected for display according to adjusted *P* value. Complete simplified results for all three ontologies are provided in **Supplementary Table 8**.

**Cross-cohort validation of sex-associated proteins**

To orthogonally validate the Plasmix M/F contrast and define externally supported sex-associated proteins, we analyzed summary statistics derived from two independent population-scale studies. Effect estimates ($\beta_{sex}$) from UK Biobank and Iceland were sign-inverted to represent the male-versus-female contrast used by Wellness, BAMSE and Plasmix. $P$ values were adjusted separately within each cohort using the Benjamini–Hochberg procedure.

Analytical records were mapped to canonical UniProt accessions. A cohort was counted as supporting a protein when at least one mapped assay record had FDR <0.05 and an absolute effect estimate ≥0.05. Opposing supported effects among the retained assay records resulted in classification as Conflict. Among non-conflicting proteins, Tier 1 required support in at least three cohorts with representation in both Olink and SomaScan; Tier 2 required support in at least three cohorts within one platform or in two cohorts across platforms; and Tier 3 comprised proteins supported in one or two cohorts without meeting the higher-tier criteria.

For cohort-level comparisons, multiple assay records mapping to the same UniProt accession within a cohort and platform were summarized using the median effect estimate and the minimum P value and FDR. Pairwise consistency classes for UK Biobank versus Iceland and Wellness versus BAMSE were based on cohort-level FDR <0.05 and effect direction. For comparison between the two study sources, cohort-level effects were averaged within the UK Biobank–Iceland and Wellness–BAMSE sources, respectively. Pearson and Spearman correlations were calculated separately by evidence tier.

Tier 1 proteins were subsequently compared with Plasmix M/F effects from Baseline-tier DIA and Calibrated-tier Olink and SomaScan measurements. Plasmix analytical features mapping to Tier 1 proteins were retained separately. Correlations were calculated independently within each Plasmix batch and summarized across batches by their median and range. For the heatmap, Tier 1 proteins represented on at least two Plasmix platforms were ranked by their maximum absolute Plasmix M/F effect, and the leading 50 proteins were displayed.

**Population-scale comparison of the Plasmix M/F effect-size range**

Plasmix Olink Explore HT measurements were compared with individual-level UK Biobank Olink Explore 3072 NPX data and external Olink Explore HT cohort summaries. Targets were matched across panel generations using the Olink mapping ledger and canonical UniProt accessions. UK Biobank M/F effects were estimated from participant-level NPX values using limma with a male-versus-female group design. Plasmix effects were obtained from the two Olink Explore HT NPX batches and summarized by the median after requiring representation in both batches and concordant effect direction.

Pearson and Spearman correlations between UK Biobank and Plasmix effects were calculated for all matched targets and separately for Tier 1 proteins. Excess-shift proteins were defined as matched targets satisfying both of the following conditions:

$$| \log_2(M/F)_{\mathrm{UKB}} | \leq \log_2(1.05)$$

and

$$| \log_2(M/F)_{\text{Plasmix}} | \geq 1$$

Finite-cohort resampling used the same matched target set. In each of 5,000 iterations, 54 male and 51 female UK Biobank participants were sampled without replacement. Direct resampling calculated the difference between the male and female mean NPX values. Pseudo-pool resampling first converted NPX to the linear scale, averaged values within each sex and then returned the pooled values to the $\log_2$ scale. The central 95% width was defined as the difference between the 97.5th and 2.5th percentiles of the protein-level effects.

Gene Ontology over-representation analysis of excess-shift proteins was performed for Biological Process, Cellular Component and Molecular Function using the complete matched UK Biobank–Plasmix target set as the background. Redundant terms were simplified at a semantic-similarity cutoff of 0.65.

### Cross-platform variance harmonization

#### Principal variance component analysis

To systematically decouple technical artifacts from biological variance, we performed the principal variance component analysis (PVCA) utilizing the imputed global datasets. To ensure an unbiased evaluation of global variance distributions, the analysis was restricted to comprehensive high-throughput platforms (Olink Explore HT, SomaScan, and DIA-MS), while targeted panels were excluded. The data input uniformly utilized the Baseline tier for DIA-MS and the Calibrated tier for Olink and SomaScan workflows, establishing a standardized multi-platform analytical window.

PCA was performed with unit-variance scaling, and the top principal components explaining 80% of the cumulative variance were extracted. Variance decomposition was executed using a Bayesian mixed-effects model via the brms package. The model incorporated five main effects (molecular recognition, data representation, pre-analytical preparation, detection instrument, and biological sample) and pairwise interactions among the four technical factors, as detailed in **Supplementary Table 9**. Weakly informative priors based on the Student's t-distribution were applied to ensure model convergence. The variance components were extracted, weighted by the eigenvalue of their corresponding principal component, and aggregated to quantify the global proportion of variance attributed to each technical or biological modality.

#### Sample-to-reference ratio and evaluation

Input data across all technology platforms were anchored at the Baseline tier. Within each analytical batch, Sample P served as the primary intra-batch reference, whereas Sample N was evaluated as an alternative anchor in the reference-choice analysis. For each feature and batch, the SRR transformation subtracted the arithmetic mean of the finite log2 measurements of the selected reference from every measurement.

The performance shift driven by SRR harmonization was evaluated using a curated subset of high-quality batches. At the sample level, PCA and signal-to-noise ratio (SNR) assessments were conducted to track clustering trajectories. At the feature level, technical precision was quantified by calculating the intraclass correlation coefficient (ICC) for technical replicates using a two-way mixed-effects model based on absolute agreement. Additionally, cross-platform fold-change concordance was evaluated by computing empirical log2 fold-changes directly from unharmonized baseline intensities. Features were classified into pairwise consensus signatures if verified as differentially expressed proteins (DEPs) in two distinct platforms, and triple consensus signatures if demonstrating agreement across all three evaluated modalities.

To evaluate global integration robustness across the comprehensive dataset without batch-level restrictions, the Symmetric Mean Absolute Percentage Error (sMAPE) was employed. Unlike variance-dependent metrics such as ICC, sMAPE provides a direct evaluation of absolute quantitative discrepancies without relying on intrinsic biological variance. For each feature within a unique pairwise batch combination, point-to-point relative percentage deviations were calculated exclusively across the non-reference study fractions (M, Y, X, and F). The integration error for any specific batch pairing was defined as the mean of these symmetric absolute deviations, providing a normalized error profile independent of directional bias.

### Quantitative response distortion

#### Empirical response-envelope modeling

Empirical response envelopes were parameterized at both the batch and platform levels using Baseline-tier measurements. Platform-native Baseline-tier representations were retained: ExtNPX for Olink, HybNorm for SomaScan, ReadoutNorm for IPP and LFQ intensities for DIA-MS. All measurements were converted from the $\log_2$ scale to the linear intensity scale before calculation. To maintain a comparable analytical scope across the response-envelope, perturbation and physicochemical analyses, Olink analyses in Fig. 4 were restricted to the Explore HT batches. The lower-plex Explore 384 batches were excluded because their limited target coverage substantially reduced the cross-platform shared target space.

Within each batch, technical replicates were summarized for each feature–sample combination by the mean linear intensity and replicate coefficient of variation (CV). The resulting observations from M, Y, P, X, F and N were pooled across samples and partitioned using intensity-percentile boundaries at 0, 0.1, 1, 2, …, 99, 99.9 and 100%, yielding 102 consecutive intensity intervals. For each interval, we calculated the number of feature–sample observations, mean linear intensity, mean HPA-derived $\log_{10}$ blood concentration, median replicate CV and proportion of observations with an HPA concentration estimate. Platform-level interval summaries were obtained by averaging the corresponding interval statistics across batches.

The relationship between estimated blood abundance and measured signal was represented by a four-parameter logistic response envelope:

$$S(x) = A + \frac{D - A}{1 + (C/x)^B},$$

where $S(x)$ is the expected linear signal at abundance $x$, and $A$ and $D$ are the mean intensities of the lowest and highest intervals, respectively. The $\log_{10}$ abundance midpoint was defined as the observation-count-weighted mean of the interval-level mean $\log_{10}$ HPA concentrations for the two intervals surrounding the median intensity boundary; $C$ was obtained by conversion to the linear abundance scale.

The signal positions corresponding to 1% and 99% of the empirical $A$–$D$ response range were defined as

$$S_1 = A + 0.01(D - A)$$

and

$$S_{99} = A + 0.99(D - A).$$

Their corresponding $\log_{10}$ HPA abundances, $h_1$ and $h_{99}$, were obtained by interpolation. The response slope was then calculated as

$$B = \frac{2\log_{10}(99)}{h_{99} - h_1}.$$

The response envelopes were therefore parameterized from empirical intensity and abundance anchors rather than estimated by unconstrained nonlinear regression.

Background burden was calculated as

$$b = 100 \times \frac{D_{\mathrm{bg}} - A_{\mathrm{bg}}}{A_{\mathrm{bg}}},$$

where $A_{\mathrm{bg}}$ and $D_{\mathrm{bg}}$ are the mean intensities of the second-lowest and second-highest intervals, respectively. Inner-tail intervals were used to reduce sensitivity to the extreme intensity endpoints. Background burden was treated as an empirical signal-domain susceptibility measure rather than a direct measurement of nonspecific binding.

Lower-, middle- and upper-range error components were estimated as the median replicate CVs within the lower 20%, middle 60% and upper 20% of the intensity distribution, respectively. At each abundance, the modeled relative variation was

$$\mathrm{CV}_{\mathrm{total}}(x) = \sqrt{\left(\frac{A\,\mathrm{CV}_{\mathrm{low}}}{S(x)}\right)^2 + \mathrm{CV}_{\mathrm{mid}}^2 + u(x)^6\mathrm{CV}_{\mathrm{high}}^2},$$

where $u(x) = (S(x) - A)/(D - A)$ denotes the relative position within the response range. Three-standard-deviation variability envelopes were generated on the $\log_{10}$ signal scale by adding and subtracting $\log_{10}[1 + 3\mathrm{CV}_{\mathrm{total}}(x)]$ from $\log_{10} S(x)$. For visualization, stochastic trajectories were generated by adding zero-mean Gaussian deviations with a standard deviation equal to 0.3 times the local envelope width.

The lower response bound was defined as $A + 3A\mathrm{CV}_{\mathrm{low}}$, and the upper response bound as $D - 3D\sqrt{\mathrm{CV}_{\mathrm{mid}}^2 + \mathrm{CV}_{\mathrm{high}}^2}$. The corresponding abundance bounds were obtained by inversion of the response envelope, and their difference on the $\log_{10}$ scale was reported as the apparent abundance span.

**Simulation of ratio distortion**

Two complementary simulations were performed using the platform-level response-envelope parameters. For the within-platform simulation, the denominator abundance was positioned one $\log_{10}$ unit below the platform-specific response midpoint, $x_1 = C/10$. Numerator abundances were defined as $2x_1$or $0.5x_1$, corresponding to input ratios of 2 and 0.5, respectively, and their expected signals were calculated from the parameterized response envelope.

A dimensionless added matrix load $L$was evaluated at zero and at 100 logarithmically spaced values from $10^{-3}$to $10^3$. For platform $p$, the mean signal-domain interference was defined as

$$\mu_p(L) = L \times \frac{b_p}{100} \times (D_p - A_p),$$

where $b_p$ is the platform-level background burden expressed as a percentage. Independent non-negative interference terms for the numerator and denominator were generated as the absolute values of Gaussian draws with mean $\mu_p(L)$and standard deviation $0.30\mu_p(L)$. The observed ratio was calculated as

$$R_{\mathrm{obs}} = \log_2 \left[\frac{S(x_2) + \varepsilon_2}{S(x_1) + \varepsilon_1}\right].$$

For each platform, input ratio and matrix-load combination, 1,000 simulations were performed. The mean and the 5th and 95th percentiles of the observed $\log_2$ ratios were reported.

For the cross-platform simulation, seven input $\log_2$ abundance differences from −3 to 3 were crossed with matrix loads of 0, 1, 3, 10 and 30. The denominator abundance was again fixed at $C/10$, and the numerator abundance was calculated as $x_1 2^{\Delta}$, where $\Delta$is the input $\log_2$ ratio. For each platform and matrix load, the deterministic interference value $\mu_p(L)$was added equally to the numerator and denominator signals before calculation of the observed $\log_2$ ratio.

The resulting SomaScan and Olink ratios were plotted on the horizontal and vertical axes, respectively, to generate the cross-platform deformation grid.

**Assessment of matrix-perturbed ratio deviations**

Matrix-perturbed ratio deviations were evaluated at the Baseline tier using features that satisfied the platform-specific batch-level detection criteria described above. Measurements were converted from the $\log_2$ scale to the linear scale before calculation. For each batch and analytical feature, technical replicates were summarized by their median linear intensities for the male pool, female pool and NIST SRM 1950 plasma, denoted $M_i$, $F_i$ and $N_i$, respectively, for feature–batch observation $i$. The native biological ratio was calculated as $r_{\text{native},i} = \log_2(M_i/F_i)$.

For affinity-based assays, buffer-blank subtraction was performed by mapping the batch- and feature-specific median raw blank signal into the Baseline-tier intensity space. Raw and Baseline-tier measurements from the same analytical well were matched using the well identifier. For sample $s \in \{M, F\}$ and replicate well $j$, the blank-subtracted Baseline-tier intensity was calculated as

$$\tilde{I}_{sij}^{\text{BLK}} = \max\left(I_{sij}^{\text{raw}}, -I_i^{\text{BLK,raw}}\ 10^{-5}\right)\frac{I_{sij}^{\text{base}}}{I_{sij}^{\text{raw}}},$$

where $I_{sij}^{\text{raw}}$ and $I_{sij}^{\text{base}}$ are the raw and Baseline-tier linear measurements from the same analytical well, and $I_i^{\text{BLK,raw}}$ is the median raw blank measurement for the corresponding feature–batch observation. A well was retained only when $I_{sij}^{\text{raw}} > 1.1 I_i^{\text{BLK,raw}}$, and all M and F replicates were required to satisfy this criterion. The corrected well-level values were summarized by the median within each sample, yielding $\tilde{M}_i^{\text{BLK}}$ and $\tilde{F}_i^{\text{BLK}}$. The blank-subtracted ratio was then calculated as $r_{\text{BLK},i} = \log_2\left(\tilde{M}_i^{\text{BLK}}/\tilde{F}_i^{\text{BLK}}\right)$. DIA-MS was not evaluated under blank subtraction because corresponding blank measurements were unavailable.

For the heterologous-plasma perturbation, observations were retained only when $M_i > 1.1N_i$ and $F_i > 1.1N_i$. The NIST-plasma-subtracted ratio was calculated as $r_{N,i} = \log_2[(M_i - N_i)/(F_i - N_i)]$.

Distribution-level distortion was quantified using the central 95% width of the corresponding ratio distribution:

$$\Delta_{95}(r) = Q_{0.975}(r) - Q_{0.025}(r).$$

The distribution-level expansion factor was subsequently defined as

$$E_{\Delta 95} = \frac{\Delta_{95}\left(r_{\text{perturbed}}\right)}{\Delta_{95}\left(r_{\text{native}}\right)}.$$

Here, $E_{\Delta 95}$ denotes the distribution-level expansion factor reported in **Fig. 4e**. Batch-level widths were calculated separately within each analytical batch, whereas platform-level widths were calculated after pooling all eligible feature–batch observations within the corresponding platform. For affinity-based assays, BLK- and NIST-plasma-perturbed distributions were evaluated using the matched set of observations satisfying the validity criteria for both perturbations. DIA summaries included observations satisfying the NIST-plasma criterion.

For feature-level predictive modeling, susceptibility to NIST-plasma perturbation was defined separately from the distribution-level expansion factor. Native ratios with $| r_{\text{native},i} |< 0.01$were excluded because division by near-zero ratios produced unstable estimates. Relative expansion for each remaining feature–batch observation was defined as

$$E_i = \frac{r_{N,i}}{r_{\text{native},i}}.$$

Only finite, positive values of $E_i$were retained. The response used for machine-learning analysis was then defined as

$$y_i = \log_2(E_i).$$

The models therefore predicted $\log_2$-transformed relative expansion rather than the absolute magnitude of ratio compression.

### Physicochemical susceptibility

#### Physicochemical feature preparation

A panel of 30 candidate structural and physicochemical properties was compiled for each canonical UniProt target using protein-sequence annotations, structural information and reference-abundance data, with computational definitions and sources detailed in **Supplementary Table 10**. Pairwise Spearman correlations and interpretability-guided redundancy assessment were used to reduce the candidate panel. Near-collinear properties were represented by the more directly interpretable descriptor, whereas strongly size-dependent absolute surface measurements were replaced by size-normalized ratios. These steps yielded 22 non-redundant properties spanning structure, surface characteristics, charge, disorder, secretory features and circulating abundance.

Informative missingness was evaluated separately within each platform. For every property containing missing values, the distributions of untransformed relative expansion $E_i$were compared between observations with and without an available property value using a two-sided Wilcoxon rank-sum test. Cliff's delta was calculated as an effect-size measure, and P values were adjusted within each platform using the Benjamini–Hochberg procedure. Missingness was considered non-trivially associated with expansion when the adjusted $P < 0.05$ and $| \delta |\geq 0.15$. Circulating abundance, alpha-helix fraction and beta-sheet fraction met these criteria and were excluded, leaving 19 properties for predictive modeling.

#### Physicochemical modeling of perturbation susceptibility

Random Forest, XGBoost and LightGBM regressors were trained independently for DIA, Olink and SOMAmer-based assays. Two target scopes were evaluated. The platform-wide analysis included all eligible feature–batch observations within each platform. The platform-shared analysis was restricted to observations whose canonical UniProt targets were represented in all three platforms; individual analytical features and batches remained separate modeling observations. A Gaussian random-noise variable generated independently for each platform and target scope was included as a negative-control predictor.

Random Forest models were implemented using randomForest with 500 trees, $mtry = 5$ and a terminal-node size of 5. Missing continuous predictors were imputed using the training-fold median, and binary predictors using the training-fold majority state; the resulting values were applied unchanged to the corresponding test fold. XGBoost models were implemented using xgboost with 100 boosting rounds, learning rate 0.05, maximum depth 3, minimum child weight 3, column subsampling of 0.8 and row subsampling of 0.8. LightGBM models were implemented using lightgbm with 100 boosting rounds, learning rate 0.05, maximum depth 3, seven leaves, a minimum of five observations per leaf, feature fraction 0.8 and bagging fraction 0.8. XGBoost and LightGBM retained missing predictor values for native algorithmic handling.

Predictive performance was assessed using 50 repeats of row-wise fivefold cross-validation. Within each repeat, feature–batch observations were randomly assigned to balanced folds, and identical fold assignments were used for all three algorithms. Predictions from the five held-out folds were pooled, and $R^2$ was calculated for each repeat as

$$R^2 = 100\left[1 - \frac{\sum_i (y_i - \hat{y}_i)^2}{\sum_i (y_i - \bar{y})^2}\right].$$

Results were summarized as the mean ± s.d. across repeats. Because folds were assigned at the feature–batch-observation level, this analysis evaluated prediction across observed feature–batch contexts rather than extrapolation to wholly unseen proteins.

Final models were subsequently fitted to all eligible observations for variable-importance and response-profile analyses; cross-validated predictions were used exclusively for performance assessment. Random Forest importance was quantified using percentage increase in mean squared error, whereas XGBoost and LightGBM importance was quantified using Gain. Negative importance values were set to zero, and importance was normalized separately within each platform, target scope and algorithm so that the largest value equalled 100:

$$VI_j^{\mathrm{rel}} = 100\frac{\max(VI_j, 0)}{\max_k\{\max(VI_k, 0)\}}.$$

The consensus importance score was the arithmetic mean of the three algorithm-specific relative importance values. It was used only as a relative ranking measure and was not scaled by model $R^2$. Properties displayed in **Fig. 4g** comprised the union of the five highest-ranked platform-wide properties across the three platforms; platform-shared importance was shown for the same property set.

Two-dimensional partial-dependence surfaces were estimated from the platform-shared Random Forest models using the pdp package. Property pairs were selected from the two highest platform-wide consensus ranks for each platform: Delta pI and low-pLDDT fraction for SOMAmer-based assays, glycosylation density and sequence instability for Olink, and glycosylation density and net charge for DIA. Surfaces were evaluated on grids with a resolution of 30 and restricted to the convex hull of the observed predictor space. At each grid value, that predictor was replaced for all training observations while all other predictors were retained, predictions were averaged on the fitted $\log_2$ scale, and the resulting mean was transformed as $2^{\hat{y}}$.

**Physicochemical association and response-profile analyses**

Pairwise Spearman correlations among the 30 numerical candidate properties were calculated across canonical UniProt entries using pairwise complete observations. Properties were ordered by Ward.D2 hierarchical clustering using $1-|\rho|$as the distance measure. Two-sided Spearman correlation tests were performed without multiple-testing adjustment for this descriptive analysis, and correlations with $P > 0.05$were marked in **Extended Data Fig. 6a**.

To test property differences across the observed expansion distribution, feature–batch observations were ranked within each platform and target scope by $E_i$. At tail thresholds from 5% to 50% in 5-percentage-point increments, the upper and lower tails were compared for each of the 22 redundancy-filtered properties using two-sided Wilcoxon rank-sum tests. A comparison was performed only when at least ten non-missing observations were available in each tail. Unadjusted *P* values were retained for the threshold-scan matrix; **Extended Data Fig. 6b** displays the platform-wide analysis.

One-dimensional partial-dependence profiles were calculated from the final Random Forest models for both target scopes. Predictor grids comprised up to 30 quantile-based points spanning the 2nd to 98th percentiles of the observed distribution. At each grid value, that predictor was replaced for all training observations while all other predictors were retained, predictions were averaged on the fitted $\log_2$ scale, and the resulting mean was transformed as $2^{\hat{y}}$. PDPs were therefore displayed on the original relative-expansion scale.

Accumulated-local-effect profiles were computed on the fitted $\log_2$ scale using 20 quantile-defined intervals. Within each interval, the predictor was replaced by its lower and upper boundary values, local prediction differences were averaged, and these differences were accumulated and centred using interval counts as weights. Binary properties were evaluated by the direct difference between their two states and centred symmetrically around zero. Only properties displayed in **Fig. 4g** were included in **Extended Data Fig. 6c,d**. For visualization alone, isolated predictor values lying below $Q_1 - 2$ IQR or above $Q_3 + 2$ IQRwere omitted from the plotted x-axis range; the calculations and exported source data retained the complete predictor range.

### Integration benchmark

#### Simulation of batch-to-batch overlap designs

All 66 batch pairs were evaluated under three study-sample overlap designs, producing 198 batch-pair–design tasks. In the Balanced design, both batches contained M, Y, P, X, F and N, and quantitative agreement was evaluated across the four non-reference study samples M, Y, X and F. In the Partial design, batch 1 contained M, Y, P, X and N, whereas batch 2 contained Y, P, X, F and N; quantitative agreement was therefore evaluated across the shared study samples Y and X. In the Confounded design, batch 1 contained M, Y, P and N and batch 2 contained X, F, P and N, leaving no shared study sample and retaining only P and N as potential cross-batch bridges.

For Confounded designs, P-anchored methods were quantitatively validated using the independently measured non-anchor sample N, whereas N-anchored methods were validated using P. Native and RF-BECA methods did not contain an explicit reference anchor and were therefore evaluated in two parallel tracks: an N validation track corresponding to the P-anchor comparison and a P validation track corresponding to the N-anchor comparison. For method-level pair summaries, the two track-specific rates were averaged with equal weight.

#### Feature eligibility and integration-success criteria

Only analytical features with an unambiguous canonical protein assignment were considered. For each batch pair, up to three pre-specified technical replicates per included sample group were retained according to the fixed sample-metadata order. Initial eligibility was determined from the primary final representation. The M and F endpoint means had to be finite and supported by at least two replicate measurements in both batches, and a finite feature-specific MAPD estimate had to be available in both batches. A feature was retained when the absolute M/F difference exceeded three times the corresponding MAPD in at least one of the two batches. The retained feature also had to be complete across all design-available samples in both the Baseline and final analysis branches. No missing-value imputation was performed in the integration benchmark. Detection status and native titration monotonicity were not used as feature-level pre-filtering criteria.

All corrected measurements were converted to the linear scale before evaluation. Quantitative agreement was determined using the design-specific sMAPE calculation described above. Expected-response retention was evaluated from the pooled corrected M, Y, X and F measurements; P was excluded because it could serve as the normalization anchor, and N was not part of the predefined Plasmix titration series. Group centres were calculated as arithmetic means and required at least two finite replicate measurements. M and F defined the endpoint coordinates of 0 and 1, and Y and X had nominal TRC positions of 0.25 and 0.75, respectively. Both intermediate TRCs had to be finite, their mean absolute deviation from the nominal positions had to be <0.25, and the corrected M–Y–X–F trajectory had to satisfy the strict titration-monotonicity criterion described above. A feature was classified as harmonized only when it simultaneously achieved sMAPE ≤ 30% and retained the expected response.

For each method, batch pair, design and validation track, quantitative-agreement, expected-response and harmonized rates were calculated as the proportions of eligible features satisfying the respective criteria. Non-finite evaluations were counted as failures. Where two Confounded validation tracks were defined for Native and RF-BECA methods, the pair-level rates reported in Fig. 6b and Supplementary Table 14 were calculated by equally averaging the two track-specific rates.

**Method- and strategy-level integration summaries**

For Fig. 6b, the rate for each individual method within a platform-pair scenario and overlap design was summarized as the median across batch pairs, thereby assigning equal weight to every pair. Strategy-level trajectories were subsequently calculated as the median of the corresponding method-level values. P- and N-anchored SRR strategies and their SRR+RF-BECA combinations were retained as distinct strategy classes in this panel.

For Fig. 6c and Supplementary Table 15, method-level results were first collapsed into five strategy classes: Native, RF-BECA, RI-BECA, SRR and SRR+RF-BECA. A strategy was considered successful for a feature–batch-pair combination when at least one constituent method satisfied the overall harmonization criterion. For Native and RF-BECA methods in the Confounded design, success in either validation track was sufficient for the feature-level strategy call used in this mutually exclusive analysis. Features were classified as Shared success when at least one SRR-enabled strategy and at least one non-SRR strategy succeeded. The remaining features were assigned hierarchically to SRR, SRR+RF-BECA, RF-BECA, RI-BECA, Native or Unresolved, in that order. Category proportions were calculated separately within each batch pair and were then averaged across batch pairs within each scenario and design.

For Fig. 6d, consensus integration success was evaluated using the Balanced design. Within every feature–batch-pair combination, the success rate was first calculated as the proportion of the 20 pipelines satisfying the overall harmonization criterion. The feature-level consensus success rate for each platform-pair scenario was then defined as the median of these pair-specific rates across all evaluable batch pairs. Features were ranked in descending order of consensus success. A separate SRR(P) diagnosis was derived from the Balanced-design feature-level results. Within each platform-pair scenario, a feature was classified as SRR(P) Pass when SRR(P) satisfied the overall harmonization criterion in at least one evaluable batch pair, corresponding to an SRR(P) success rate greater than zero, and as Fail otherwise. Features with a consensus success rate of zero were designated unresolved across all evaluated pipelines.

**Reference-choice and background-subtracted SRR analyses**

Extended Data Fig. 8a compared P- and N-anchored SRR under the Balanced design. For each reference, batch pair and platform-pair scenario, quantitative-agreement, expected-response and harmonized rates were taken from the same feature-level analysis used for Fig. 6. Displayed scenario-level values were the medians of the corresponding pair-level rates.

For the blank-subtracted SRR analysis in Extended Data Fig. 8c, affinity-platform Raw readout and Baseline-tier measurements from the same analytical well were matched. For every feature and batch, the median Raw readout of the buffer blanks was calculated. For wells with finite positive Raw readouts, the well-specific mapping factor from Raw readout to the Baseline tier was calculated as the Baseline measurement divided by the Raw measurement. The background-subtracted Baseline value was then calculated as the larger of the blank-subtracted Raw measurement and $10^{-5}$, multiplied by the well-specific mapping factor. DIA-MS measurements were retained unchanged because corresponding buffer-blank measurements were unavailable.

For each feature and batch, the arithmetic mean of the background-subtracted P or N measurements was used as the reference denominator, and all analytical samples were expressed as linear sample-to-reference ratios. The analysis used the same Balanced-design feature eligibility sets as the corresponding standard SRR(P) and SRR(N) analyses. Intra-DIA comparisons were omitted, whereas DIA measurements were retained unchanged in cross-platform comparisons involving an affinity platform. Quantitative agreement was evaluated across M, Y, X and F, and expected-response and harmonized classifications followed the same criteria used in the primary integration benchmark.

**Abundance-associated integration outcomes**

For Extended Data Fig. 9, the Balanced-design consensus success rate described above was used to define scenario-specific outcome groups. High-consensus proteins were those with a consensus success rate greater than or equal to the 75th percentile within the corresponding platform-pair scenario. Failed proteins were those with a consensus success rate of zero, and proteins with intermediate rates were excluded from this comparison.

Native measurement ranks were calculated from the same primary final representation used in the integration analysis. Within each analytical batch, the median native measurement was calculated for every feature and converted to a percentile rank. For each batch pair, the lower of the two batch-specific percentile ranks was retained to represent the lower-measurement side of the comparison. The feature-level pair-low-side native measurement rank was the median of this value across all evaluable batch pairs within the scenario. HPA-derived circulating concentrations were mapped through canonical UniProt accessions and expressed as $\log_{10}$ pg/mL. The distributions of the native measurement rank and HPA concentration were compared descriptively between high-consensus and failed proteins.

**Physicochemical associations with integration outcomes**

Physicochemical associations with integration success in Fig. 6e and Supplementary Table 16 were evaluated using the same scenario-specific high-consensus and failed groups. Proteins with intermediate consensus success rates were excluded. Consensus classifications were joined to the curated physicochemical matrix through canonical UniProt accessions, and all properties retained in the physicochemical annotation dictionary were evaluated.

For each platform-pair scenario and physicochemical property, finite property values were compared between high-consensus and failed proteins when at least five proteins were available in each group. Distributional differences were evaluated using a two-sided Wilcoxon rank-sum test, and Cliff's delta was calculated as a non-parametric effect-size measure. Positive Cliff's delta values indicated higher property values among high-consensus proteins, whereas negative values indicated higher values among failed proteins. Figure annotations were based on nominal $P$-value thresholds of 0.05, 0.01 and 0.001. The figure displayed properties that were nominally significant in at least one scenario, while Supplementary Table 16 reports all evaluated properties together with Benjamini–Hochberg-adjusted $P$ values calculated separately within each platform-pair scenario.
